# A review of fish diversity in Mediterranean seagrass habitats, with a focus on functional traits

**DOI:** 10.1101/2023.09.23.559097

**Authors:** A. Lattanzi, B. Bellisario, R. Cimmaruta

## Abstract

Seagrass habitats are considered essential in supporting both fish diversity and fisheries’ income. Within the Mediterranean Sea most fish species are associated to seagrasses and constitute the subject of extensive literature. However, a complete synthesis of fish species observed in different seagrass habitats is still lacking at the whole-basin Mediterranean scale, so hindering a thorough understanding of the main functional traits and mechanisms involved in determining fish diversity patterns. We performed a systematic review by implementing a semi-automated, threshold-based filtering pipeline that allowed building up a dataset concerning all fish species reported in native Mediterranean seagrasses, including specific functional traits known to be involved with the potential use of seagrasses by fish. These data allowed to carry on a narrative synthesis on fish diversity in seagrass habitats. Moreover, the functional traits analysis highlighted a significant role of trophic level, size and benthic domain in determining the distribution of fish in different seagrass habitats at whole Mediterranean basin scale. Data evidenced a pattern of segregation between large piscivorous, pelagic predators and smaller benthic prey (including juveniles) with omnivorous to herbivorous feeding habit, with aggressive predators significantly associated with *Posidonia oceanica* and *Cymodocea nodosa* beds while their putative prey tend to occupy other seagrass habitats. We identified unexpected knowledge gaps on the role of habitat heterogeneity and fish life stages in determining the presence/absence of fish in different seagrasses, for which in-depth research is needed to fully understand the links between nursery habitats and the state of exploited fish stock.

## Introduction

The Mediterranean Sea is a small, oligotrophic basin recognized as a main hotspot of biodiversity due to both the high species richness and endemicity, a well-known pattern for many animal groups, mainly crustaceans and molluscs, including fish (Coll et al. 2010). It also hosts unique and important habitats for marine diversity such as seagrass meadows, considered pivotal in maintaining fish biodiversity, being used as both foraging grounds and nursery (Jackson et al. 2015; McDevitt-Irwin et al. 2016).

Seagrasses are a unique group of flowering plants widely distributed in coastal marine environments all around the world where they are acknowledged as biodiversity hotspots (Ruiz et al. 2009). As habitat engineers, seagrasses support many ecological and economic activities including the regulation of ecological processes and the provisioning of goods and cultural values for human well-being (do Amaral Camara Lima et al. 2023). Within the Mediterranean Sea, five native species of seagrasses occur (Ruiz et al. 2009). The endemic large Neptune seagrass *Posidonia oceanica* (L.) Delile has a paramount importance in terms of ecosystem services and biodiversity along with *Cymodocea nodosa* (Ucria) Ascherson, *Zostera marina* L. and *Z. noltei* Hornemann 1832 that have a broader temperate distribution, and *Ruppia cirrhosa* (Petagna) Grande, 1918 typical of brackish lagoons and salt marshes (Duarte 2009). Two more species, *Halophila stipulacea* (Forsskål) Ascherson, 1867, and *H. decipiens* Ostenfeld, 1902 are Lessepsian and ballast introduced (Gerakaris et al. 2020).

A significant number of Mediterranean fish use seagrass habitats to forage, from herbivorous grazers to omnivores and predators that may feed upon animals living in the canopy where the shaded and colder waters provide shelter for species at different life stages (Terrados and Duarte 2000; Boudouresque 2004). Seagrass beds are also important nursery habitats for several species of fish that here spawn or settle as juveniles, either spending their entire life cycle in the meadows or moving to other ecosystems as they mature (Ren et al. 2022). For these reasons, seagrasses are considered of paramount importance in sustaining fisheries within the Mediterranean Sea and all around the world, supporting large populations of commercially and socially important fish species (Unsworth and Cullen 2010).

Within this framework, it is not surprising that Mediterranean seagrasses are widely studied and that systematic literature reviews are frequently carried out to both summarize the state of the art and identify future research directions (Romero et al. 2016; Boudouresque et al. 2021; Procaccini et al. 2023). However, the same attention has not being paid to fish species associated with Mediterranean seagrasses. Despite their relevance in terms of both biodiversity conservation and fisheries support, a thorough and up to date synthesis of fish species observed in different seagrass habitats is still lacking at the Mediterranean scale. Indeed, many studies are often carried out in restricted geographic areas, using unstandardized methods and pursuing different objectives, making it difficult to extract clear and comparable data. Another reason for this gap could be due to the high number of papers produced over the years, preventing to apply an expert opinion and making difficult to carry on exhaustive systematic literature reviews, usually performed to answer specific questions and therefore working on a reasonable number of papers to be filtered (Kastner et al. 2012). The subject has been therefore addressed in terms of either worldwide pattern using selected case studies (e.g., Unsworth et al. 2018), Mediterranean sub-basins (e.g., Koutsidis et al. 2020), or within specific contexts of Mediterranean fisheries or Marine Protected Areas management without a specific focus on seagrass substrates (e.g., Piroddi et al. 2020; Fraschetti et al. 2022). The lack of a comprehensive view of fish distribution in different seagrasses at the Mediterranean scale may also hinder a complete understanding of the role played by both functional traits and adaptive or behavioural mechanisms that may account for fish diversity patterns in seagrass habitats.

In this paper we tried to fill this gap by performing a systematic review through the implementation of a semi-automated, threshold-based filtering pipeline rooted into the approaches of the JBI methodology (Joanna Briggs Institute; Peters et al. 2020) and PRISMA statement (Preferred Reporting Items for Systematic Reviews and Meta-Analyses; Page et al. 2021). The aim was to produce exhaustive and comparable information on fish species that use native seagrass habitats across the entire Mediterranean Sea, by building up a dataset including all fish species reported in seagrasses with associated metadata alongside specific functional traits known to be involved with the potential use of seagrasses by fish. Data have then been used to: 1) carry on a narrative synthesis on fish diversity in seagrass habitats; 2) investigate which traits are significantly linked to the use of various seagrass habitats by different fish species; 3) identify key findings and knowledge gaps for future research. We are confident that our dataset will provide a sound basis for scientists and managers across many fields, from fisheries to biodiversity assessment and conservation. These data will enable to verify and extrapolate specific hypotheses on basin-wide scale, which might otherwise remain unverified due the complex nature of mechanisms structuring ecological communities, which are typically scale-dependent (Weiher et al. 2011).

## Materials and methods

### Systematic Review protocol

We followed the nine steps proposed by the JBI (Joanna Briggs Institute) methodology (Peters et al. 2020) and PRISMA statement (Preferred Reporting Items for Systematic Reviews and Meta-Analyses; Page et al. 2021), to collect all available information on the presence of fish species on different seagrasses of the Mediterranean Sea. Many additions were however implemented to the protocol, mainly regarding the screening phase, to cope with the high number of papers recovered by searching databases (see Supplementary Information).

The basic review question was to assess the current knowledge about fish marine biodiversity in native seagrasses at the Mediterranean Sea scale. Besides providing a narrative synthesis on fish diversity, we aimed at further addressing the following sub-questions: 1) do the species recorded on seagrasses use these habitats at all life stages? 2) which functional traits are linked to seagrass use made by different fish species? 3) which are the key findings and knowledge gaps for future research? The inclusion criteria for the review were developed in according to the PCC framework (Population (or participants)/Concept/Context), which is highly recommended for constructing a clear and meaningful title, review question, and inclusion criteria by applying the PRISMA flow diagram for the systematic search and review (Peters et al. 2020; Page et al. 2021).

We made some implementations to the screening phase of the PRISMA guidelines, with the aim of conducting an exhaustive survey of the academic literature to list all the fish species using seagrass habitats across different regions of the Mediterranean Sea and to build up an associated dataset. All types of documents were considered, and studies published in English, French, Spanish and Italian were included in the review. Grey literature was considered at the later stage of the backward and forward analysis because of the high duplication of data due to frequent grey reporting, followed by indexed publication of the same data. According to PRISMA guidelines, the survey was carried out by implementing a semi-automatic pipeline of three sequential steps, as summarized below. Detailed information for each step of the analysis is reported in Supplementary Information (SI).

### Step 1 – Identification - Searching the libraries

We conducted a keyword-based search within six different databases (Dimension, Mendeley, ScienceDirect, Scopus, Web of Science and Wiley Online Library, last access on June 30, 2023) by selecting 69 keywords identifying the three main fields of the research objective (Mediterranean Sea, seagrass, and fish fauna), linked by 68 *OR* and three *AND* Boolean operators (Fig. 1). No restrictions were applied for publication year.

**Figure 1.**
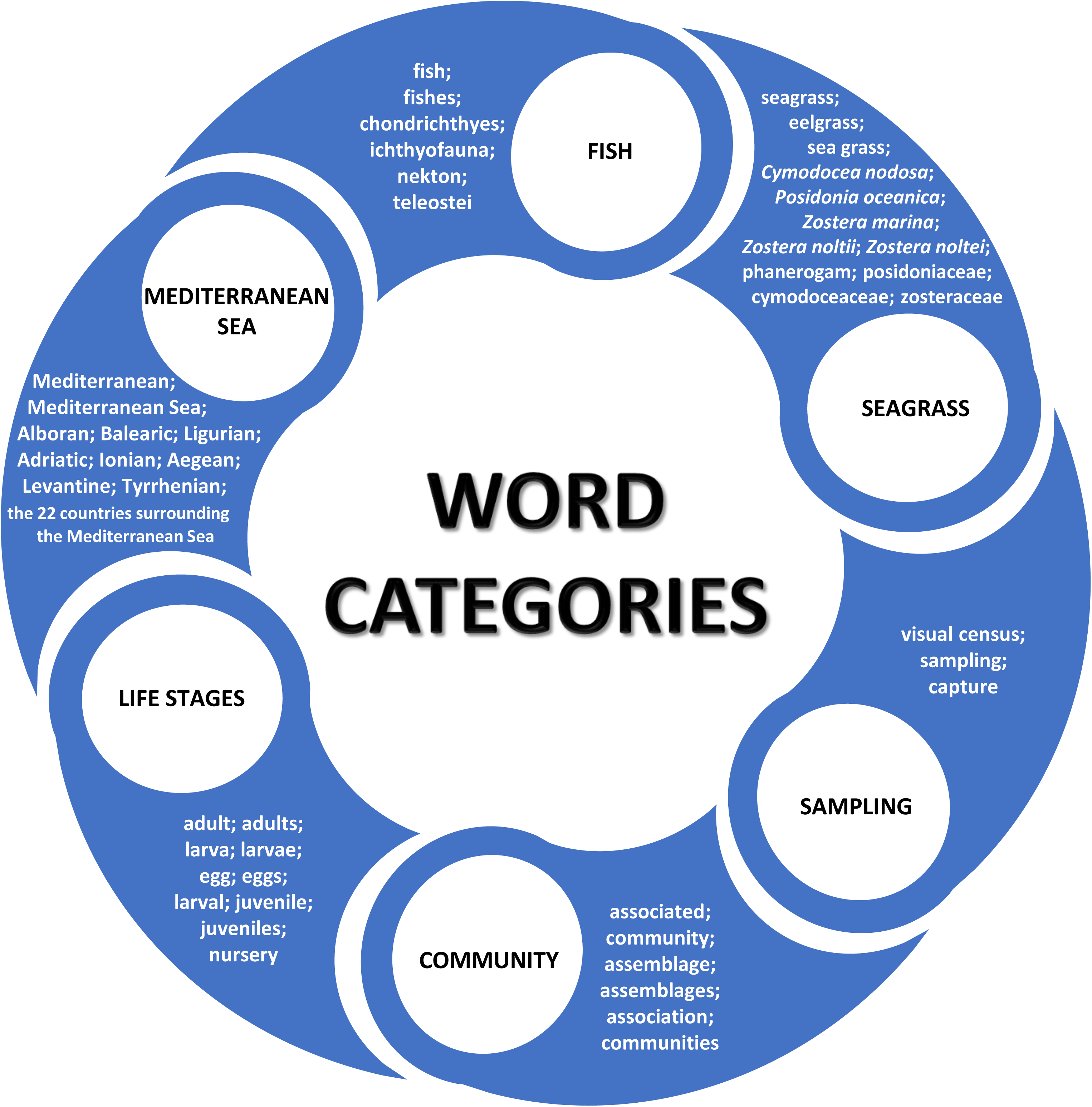
Graphical representation of the keywords used for each field of our research objective during the implementation of the filtering pipeline.

The pipeline of papers filtering is reported step by step in Fig. 2. Advanced searches were therefore carried out in each of the possible research fields (Title, Abstract, Author Keywords, Text and References) of the six bibliographic databases using a different query for each library due to both their different query limits and years of coverage (Supplementary Table S1). Results have been exported as an Excel file containing the abstract, the Authors’ keywords and the title of each paper (sub-Step 1.1 in Fig. 2), which represents the basis of subsequent searches. We removed from the paper list those records having partial searchable information (i.e., literature without abstract) and duplicate papers (sub-Step 1.2 in Fig. 2), by using both Microsoft Excel and the function *removes.duplicates* in the package ‘litsearchr’ of R (Grames et al. 2019; R Core Team 2022).

**Figure 2.**
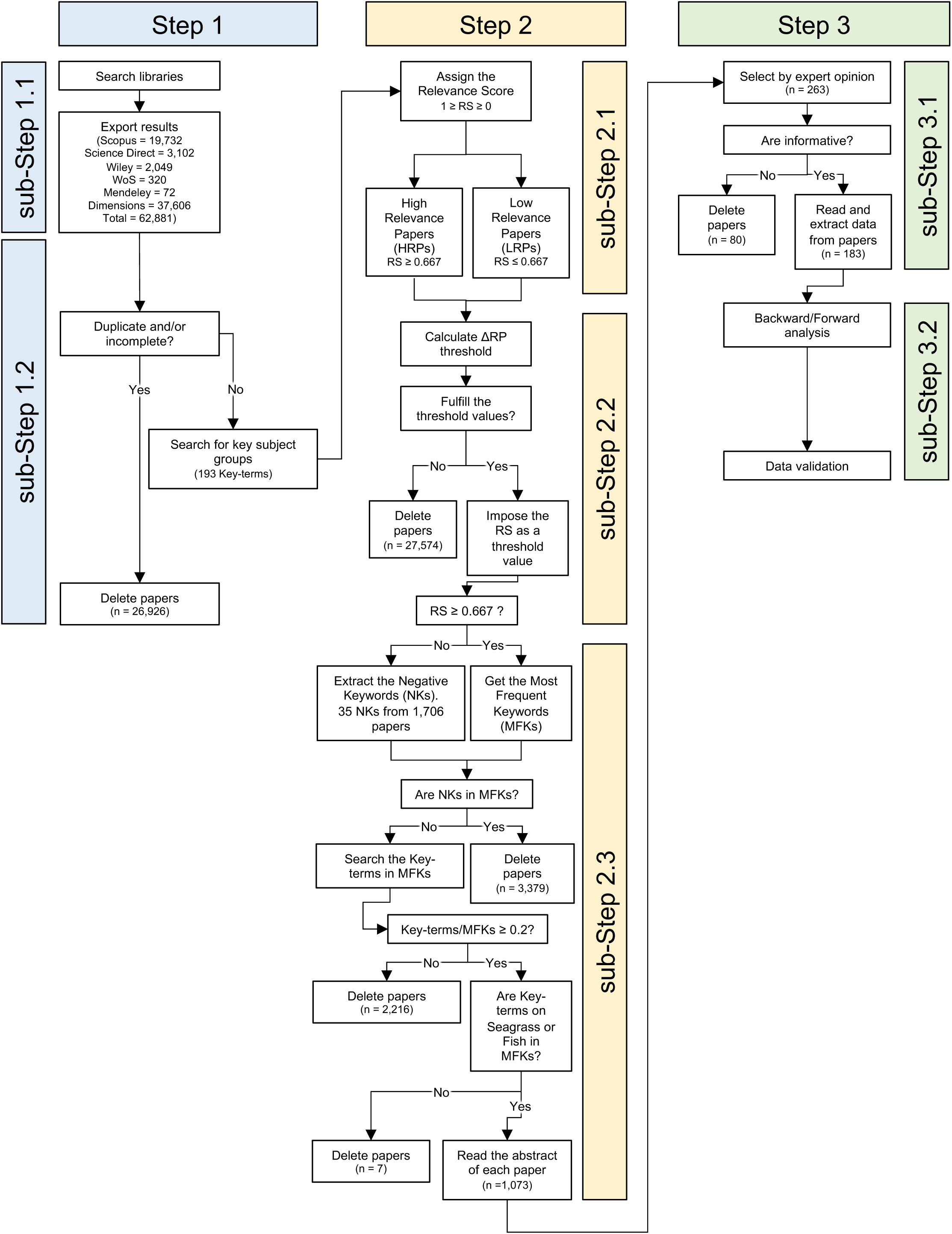
Flowchart showing each Step and sub-Step implemented in the filtering pipeline.

We set up a scoring method for the listed papers to apply several successive filters to automatically identify and discard those not containing suitable information. We therefore scored the papers by setting a higher, more specific and exhaustive set of 193 key-terms about the four key-subject groups of the research: Group S – key-terms related to the seagrasses and habitat targets; Group F – key-terms related to fish biology; Group A – key-terms related to sampling, i.e., for either the methods or fish target assemblages; Group G – key-terms related to the Mediterranean basin and geographic localities (Supplementary Table S2). The presence/absence of these key-terms was assessed using the function *str_detect* in the package ‘stringr’ of R (Wickham 2019).

### Step 2 – Screening - Scoring and filtering papers

We assigned a score to each group of key-terms to calculate a Relevance Score (RS) for each paper. This allowed identifying papers likely containing a high amount of relevant information (higher RS) and to rate the type of information, i.e., concerning seagrass habitat, and/or fish assemblages, and/or sampling methods and/or geographic locality (sub-Step 2.1 in Fig. 2). A detailed protocol is described in SI and Supplementary Table S3.

Once obtained a RS for each paper, we performed a sequential filtering to keep only those papers able to provide information suitable to build up the final dataset. Each filter was applied on the papers remaining after the previous filtration step. Filter 1 allowed excluding papers with a score lower than a threshold value based on the comparison of the representativeness of each of the four key-subject groups in papers with high *vs*. low RS (sub-Step 2.2 in Fig. 2 and Supplementary Tables S4 and S5). Filter 2 allowed dropping the papers with a low RS (sub-Step 2.2 in Fig. 2). Filter 3 was based on both the ‘Most Frequent Keywords’ (MFK, identifying recurring and relevant keywords) and ‘Negative Keywords’ (NK, whose presence indicates that the paper does not focus on Mediterranean seagrass and/or fish communities although mentioning them, for instance for comparison purposes). This last filter allowed removing: i) all papers containing one or more NK within their MFK; ii) all papers with a low key-terms/MFK ratio; iii) all papers that do not have any keyword related to Group S (Seagrass) or F (Fish) in their MFK (sub-Step 2.3 in Fig. 2). The pipeline of filtering steps is described in detail in SI.

### Step 3 – Eligibility and inclusion – Data extraction and refining

Once obtained the short list of papers likely containing useful information, we read all the abstracts to further select by expert opinion those of interest, that is, papers providing information on fish presence in seagrass habitats across the Mediterranean Sea (sub-Step 3.1 in Fig. 2). The selection process was based on the ‘team approach’, as recommended by Levac et al. (2010).

As the selected libraries could miss studies of interest, during the full text reading we searched for useful papers absent from the six libraries but listed in their references by carrying on a backward search based on the Reference papers and, similarly, forward research with the aim of integrating as much data as possible in case of partial data in the papers within our list (sub-Step 3.2 in Fig. 2).

To obtain complete and comparable data, we carried out a manual extraction of data concerning each recorded fish species, along with the source reference paper. The fish species list was finally refined by discarding singletons, thus retaining only those species recorded in at least two separate research studies (in accordance with Unsworth et al. 2014). However, among the singletons detected, we kept Lessepsian species, frequently being listed as a unique record (sub-Step 3.2 in Fig. 2).

The resulting dataset provided a full list of the fish species recorded on seagrasses by the present literature, by reporting each observation associated with: Mediterranean ecoregion, the reference FAO zone and GFCM Area, site and geographic location, season, year and sampling method, habitat composition and substrate, life stage, latitude and longitude of each location and bibliographic information (see Data Availability Statement).

### Patterns of fish diversity

To provide an overall view of fish diversity in Mediterranean seagrasses, a curve slope analysis was used to illustrate differences in the frequency of records among the 248 fish species reported in literature. To this end, we performed a piecewise regression between the number of records of listed species and the number of localities, comparing the goodness of fit with the equivalent linear regressions by the Akaike Information Criterion (AIC, Burnham and Anderson 1998) and best model was expressed by AIC differences, ΔAIC = AIC − min(AIC) and Akaike model weights. All computations were performed by using the ‘segmented’ package of R (Muggeo 2008).

We also performed a series of analyses to understand how fish species distribute over different seagrasses according to their life stage and which traits were mostly associated with the patterns detected. We first started by aggregating data contained in the ‘Habitat’ field of the dataset to homogenate the over 30 different types of seagrass and seagrass/seaweeds/coralligenous/*Ruppia* combinations found in the literature. Data were therefore aggregated in 10 main seagrass habitats subdivided in either pure (i.e., meadows constituted by a single seagrass) or mixed (i.e., seagrass in combination with other seagrass and/or seaweeds, coralligenous or *Ruppia*). We then reconstructed a species incidence matrix where each entry corresponded to the number of papers in which each fish species was reported in each habitat. We further arranged our data by characterizing the life stage at which each species was found in each seagrass habitat (adults, juveniles and both stages). Moreover, we used the FishMed database (Albouy et al. 2015) to extract species-specific traits associated with the potential use of seagrass by fish: i) average length; ii) vertical distribution; iii) diet type of larvae; iv) diet type of adults and v) trophic level.

A Canonical Correspondence Analysis (CCA) was performed to relate species compositional data (i.e., the presence/absence of fish on different seagrass habitats at a given life stage) with traits. Since selected traits were composed of both quantitative (i.e., average length and trophic level) and qualitative data (i.e., vertical distribution, diet type of larvae and adults), we first performed a Factorial Analysis of Mixed Data (FAMD, Pagès 2004) with the aim to reduce multicollinearity and the number of variables in a bi-dimensional space provided by the ordination. FAMD is a principal component method able to handle both continuous and categorical variables, where continuous variables are scaled to a unit variance, and categorical variables are transformed into a disjunctive data table and then scaled to balance the influence of both continuous and categorical variables in the analysis. The first two dimensions of the FAMD were then used as explanatory variables in the CCA, using ANOVA with 1,000 permutations to test for the statistical significance.

## Results

### Workflow and literature overview

At the first search performed, we obtained 62,881 results from the six bibliographic databases, including articles, book chapters, technical reports and conference papers. The resolution power of this research was low, since we reduced this number to 35,955 papers by just eliminating duplicates and publications showing incomplete information (Fig. 2).

The further filtering actions sorted out papers that did contain information on the presence of fish species on Mediterranean native seagrass by using a combination of keyword-based filters. The application of filter 1 reduced the total number of papers by 77%, excluding papers that did not meet the requirements about the minimum number of key-terms in title and abstract (27,574). Another 1,706 poorly relevant papers (filter 2, RS < 0.667) were further discarded resulting in 6,675 remaining papers. By using filter 3, we focused on the 1,073 papers possibly containing extractable data by eliminating those either containing Negative Keywords or not fitting Most Frequent Keywords thresholds (5,602). Abstracts were read to identify papers likely containing data of interest; 183 were retained and used as the base for *a posteriori* backward snowballing to further search possible papers not indexed in the databases or escaped to research strings. Only fifteen papers were added after this final check, indicating that the searching pipeline implemented in this work was efficient in recalling the scientific literature containing data usable for our research aims. Finally, 33 papers were discarded since reporting data already published elsewhere (for example for meta-analytic purposes) and then data were extracted from 165 papers reporting original, unduplicated data. The complete list of reference papers is available in the provided dataset (see Data Availability Statement).

An overall summary of the literature recovered is reported in Fig. 3, showing that most papers were published between 1982 and 2023, with an overall increasing trend over this time (Fig. 3a). Nearly 96% of papers were peer reviewed articles, and only a minor fraction was composed by technical reports, book chapters and master’s theses (Fig. 3b). Nearly half of the papers (94) reported data collected using visual census methods, while 73 were based on capture data and 16 used other methods (Table 1, Fig. 3c). Some studies (12) used both capture and visual census approaches. A large part of studies reporting the sampling season gathered their data across two to all four seasons (65%), while over a quarter focused on summer (27%) and a minority collected information during either the autumn, spring, or winter (4%, 3% and 1%, respectively) (Fig. 3d). Concerning the geographic coverage, a large majority of studies were from the Western Mediterranean Sea (116 over 165), while other ecoregions were characterized by a far lower number of studies, from 30 in the Adriatic Sea to 15 and 12 in the Eastern and Central Mediterranean, respectively (Table 1, Fig. 4).

**Figure 3.**
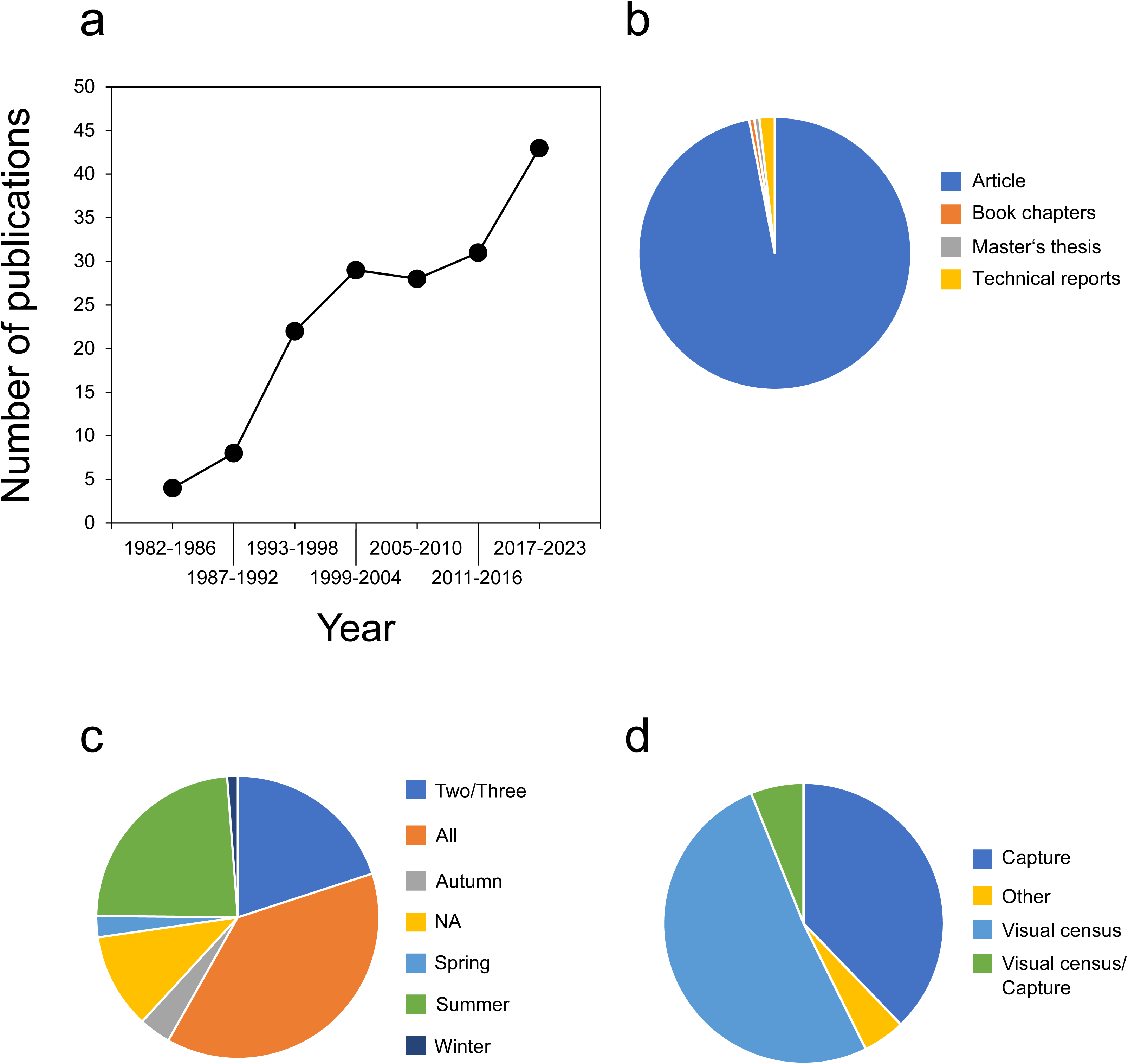
Summary of the literature overview showing: a) the number of publications, aggregated for the five-year period; b) type of publication; c) data collection seasons and d) data collection methods.

**Figure 4.**
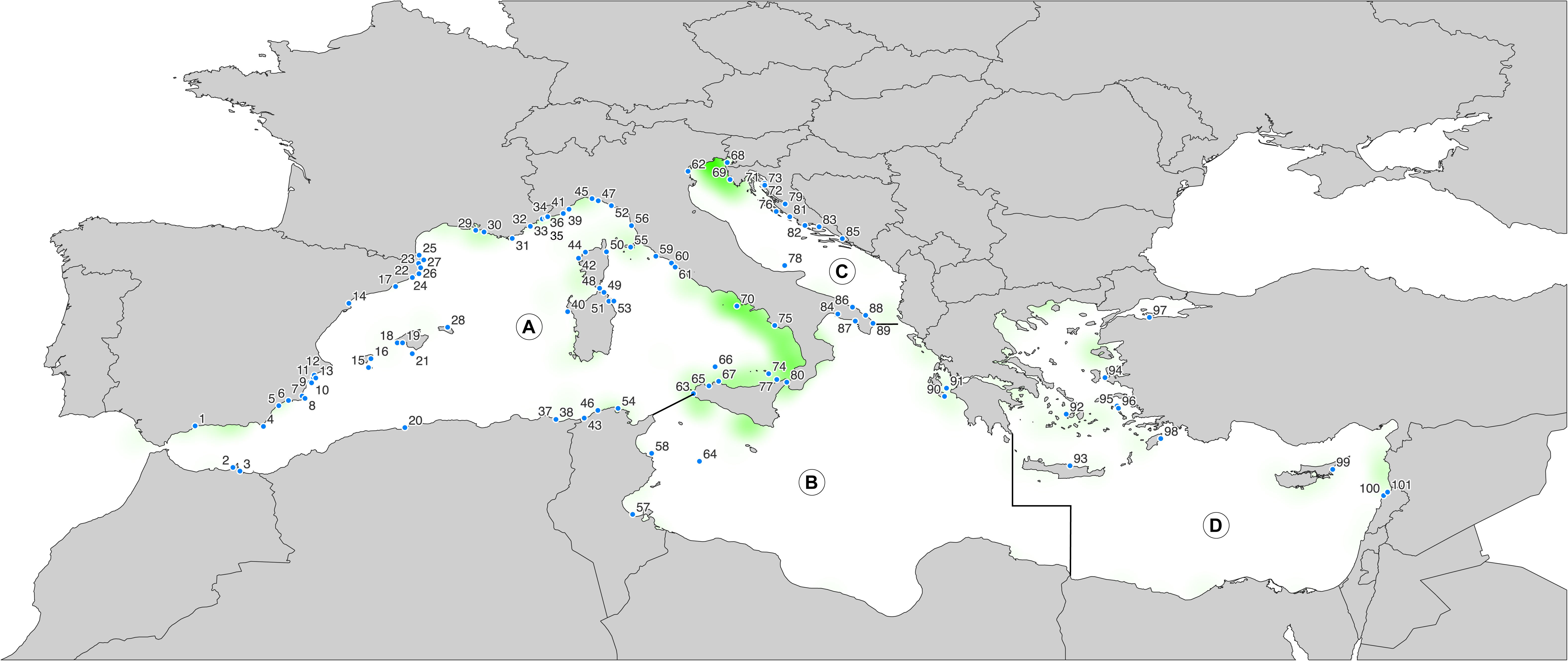
Geographic distribution of the 101 localities providing data on fish assemblages on seagrasses. A, Western Mediterranean Sea; B, Central Mediterranean Sea; C, Adriatic Sea; D, Eastern Mediterranean Sea. The density map in green was created by using data on the global distribution of native seagrasses, downloaded from https://data.unep-wcmc.org/datasets/7 and visualized as a density map in QGis. A detailed and interactive map showing all available information from our dataset can be consulted at https://brunobellisario.github.io/SeagrassFishMed.

**Table 1.** Summary table showing the total number of: i) papers, N_papers_; ii) fish species, S_tot_ and iii) exclusive fish species, S_exclusive_ with respect to different sampling methods, ecoregions, and seagrass habitats.

| | | $N_{\text{papers}}$ | $S_{\text{tot}}$ | $S_{\text{exclusive}}$ |
| --- | --- | --- | --- | --- |
| Sampling methods | Net | 73 | 225 | 29 |
|  | Visual census | 95 | 218 | 21 |
|  | Other | 17 | 76 | 1 |
| Ecoregions | Western Mediterranean | 116 | 231 | 45 |
|  | Central Mediterranean | 12 | 132 | 0 |
|  | Adriatic Sea | 30 | 162 | 4 |
|  | Eastern Mediterranean | 15 | 140 | 9 |
| Seagrass Habitat | *Posidonia oceanica* | 127 | 240 | 82 |
|  | *Cymodocea nodosa* | 28 | 146 | 2 |
|  | *Zostera marina* | 3 | 38 | 0 |
|  | *Posidonia oceanica* / *Cymodocea nodosa* | 7 | 81 | 0 |
|  | *Posidonia oceanica* / Zosteraceae | 1 | 1 | 0 |
|  | *Cymodocea nodosa* / Zosteraceae | 14 | 88 | 3 |
|  | Zosteraceae | 3 | 6 | 0 |
|  | Seagrass / seaweeds | 23 | 173 | 2 |
|  | Seagrass / Coralligenous / seaweeds | 4 | 62 | 0 |
|  | Seagrass / *Ruppia* sp. / seaweeds | 2 | 26 | 0 |

*Posidonia oceanica* meadows were the most studied habitat, with 127 papers, while 28 works reported data collected from *C. nodosa* and three regarded *Z. marina*, with another three studying mixed meadows of Zosteraceae. Mixed seagrass meadows studies were reported in 21 papers and 29 works analysed habitats characterized by a mix of seagrass with seaweeds, coralligenous or *Ruppia cirrhosa* complex (Table 1). The database regarding each study site was also visualized and disclosed on an interactive map based on Fig. 4, produced using the qgis2web plugin of QGis (QGIS Development Team 2023) and rendered via Leaflet, available at https://brunobellisario.github.io/SeagrassFishMed (hosted by GitHub, Inc.^©^ 2022).

### Fish records

A total of 248 fish species belonging to 75 families recorded from 101 localities and 10 different habitats were signalled for a total of 9,487 records. At the family level, the highest number of records concerned Labridae (1,939 records over 9,487) and Sparidae (2,316), followed by Serranidae (666), Gobiidae (611), Blenniidae (442), Syngnathidae (362), Scorpaenidae (356), Mullidae (297), Mugilidae (198) and Pomacentridae (191). Nearly all observed species were recorded by both capture (225) and visual census (218) studies, while a low number of studies carried out using other sampling methods recovered 76 species only (Table1). The greater number of species was signalled from the Western Mediterranean (231), the wider geographic area also providing the higher number of studies. A comparable number of species was recorded within the other, less investigated, ecoregions ranging between 132 (Central Mediterranean) and 162 (Adriatic Sea) (Table1).

The data extracted from literature witness that virtually all the species recorded were spotted in *P. oceanica* meadows (240 over 248) while the species pool observed in any other habitat was much more restricted, with *C. nodosa* beds hosting about 60% of the recorded species (146) and *Z. marina* only 15% (38 species) (Table 1). Habitats of mixed seagrasses were characterized by an intermediate number of recorded species; 88 in mixed meadows of *C. nodosa* with Zosteraceae and 81 in *P. oceanica*/*C. nodosa*. A low number of species was observed on mixed Zosteraceae either associated or not with *P. oceanica*. The small number of studies reported for this habitat is unlikely to be the cause of the few species recorded, since a comparable number of sources documented a higher number of species from mixed habitats of seagrass with seaweeds (Table 1).

We extracted the number of species that have been recorded as exclusive for each sampling method, ecoregion, and habitat (Table 1). Within the sampling methods, captures were able to spot 29 species unrecorded by any other method, against 21 exclusives of visual census, suggesting a certain degree of complementarity of the two methods. The number of species found exclusively in single ecoregions was high in the western Mediterranean (45) but very low in the Adriatic Sea (4). When considering the fish species observed in a single habitat, we found that about one third has been exclusively associated with *P*. *oceanica* (82 species), while other combinations of seagrass hosted a remarkably lower number of unique species; two in *C*. *nodosa* (*Apricaphanius iberus* and *Hemirhamphus far*), and three in *C. nodosa* mixed with Zosteraceae (*Hyphorhamphus picanti*, *Merlangius merlangus*, *Sygnatus taenionotus*). The pattern of fish species shared among different seagrass habitats was represented by a network that evidences the remarkable difference in the number of exclusive species observed in *P. oceanica* with respect to other seagrass habitats (Fig. 5).

**Figure 5.**
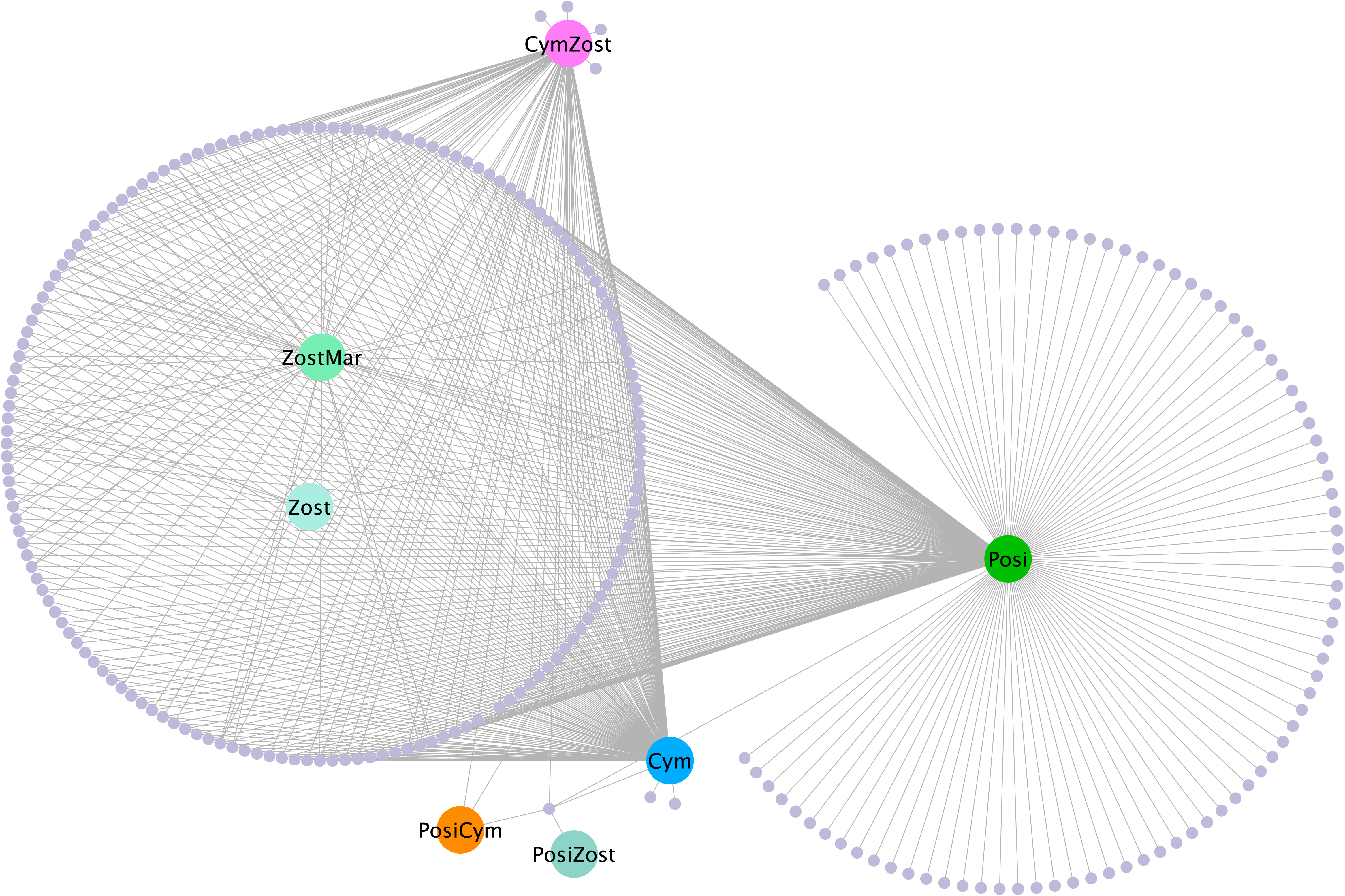
Network representing the association between fish species and native seagrasses in the Mediterranean Sea: Posi, *Posidonia oceanica*; Cym, *Cymodocea nodosa*; ZostMar, *Zostera marina*; Zost, Zosteracee; CymZost, *C*. *nodosa* mixed with Zosteracee; PosiCym, *P*. *oceanica* and *C*. *nodosa*; PosiZost, *P*. *oceanica* mixed with Zosteracee.

### Fish distribution patterns in Mediterranean seagrasses

The piecewise regression between the number of records of listed species and the number of localities improved the model fitting compared to a simple linear regression, identifying two main breakpoints (Supplementary Figure S1). Fish species can therefore be subdivided in three main frequency classes: 1) frequent and widespread species recorded many times from a high number of localities; 2) species with an intermediate frequency of records; 3) uncommon species spotted quite rarely and from a low number of localities.

We identified 23 frequent species showing between 106 and 284 records each from 55 to 85 localities. Among them, the most frequently observed were all the species belonging to the genera *Diplodus* (except *D. cervinus*) and *Symphodus* (except *S. doderleini* and *S. melops*) followed by *Mullus surmuletus* and *Coris julis*. Other species were: *Serranus scriba* and *S. cabrilla*, *Sarpa salpa*, *Chromis chromis, Boops boops*, *Oblada melanurus*, *Scorpaena porcus*, *Spondyliosoma cantharus*, *Labrus merula* and *L. viridis* and *Dentex dentex.* Another 55 species were recorded at intermediate frequencies, showing 25 to 124 records from 18 to 52 localities. Here are included for example the zebra seabream *Diplodus cervinus* and the two *Symphodus* species *S. doderleini* and *S. melops*, both sand smelts *Atherina boyeri* and *A. hepsetus*, and all the blennies belonging to the genus *Parablennius*, except the uncommon *P. zvonimiri*. The larger number of species (170) was recorded at low frequencies, being signalled 1 to 46 times from 1 to 17 localities. This long list includes species typical from coastal habitats different than seagrass, or the many pelagic species occasionally occurring on coastal seagrass meadows mainly while foraging. The 15 Lessepsian species recorded so far in seagrass habitats are also uncommon, some of them being reported by single studies (Supplementary Table S6).

One of the objectives of this review was to analyse the distribution of fish life stages across various seagrass habitats. Information on the life stage were available for 175 species in at least one study, while 73 species lacked these data and were thus excluded from further analyses. A heat-map was used to visualize at which life stage (adult, juvenile, both stages and undefined) each species was recorded in each seagrass habitat (Fig. 6a). Our findings showed that nearly all 175 species have been recorded in several studies without assessing their life stage: only 7 species were always classified from this point of view (Fig. 6a). The heat-map evidenced that a high number of species observed in *P. oceanica* meadows were reported mainly as adults, either exclusively or associated with juvenile stages. It is worth noting that nearly all species reported as adults in *P. oceanica* have been also recorded in further studies that miss the life stage. A high number of species was also recorded on *C. nodosa* but in this case the larger part of the data concerned juveniles in pure meadows and both adults and juveniles in mixed meadows of *C. nodosa* with *P. oceanica* (Fig. 6a). In the case of *C. nodosa* too, many studies did not report the life stage of the recorded specimens, leaving a knowledge gap open.

**Figure 6.**
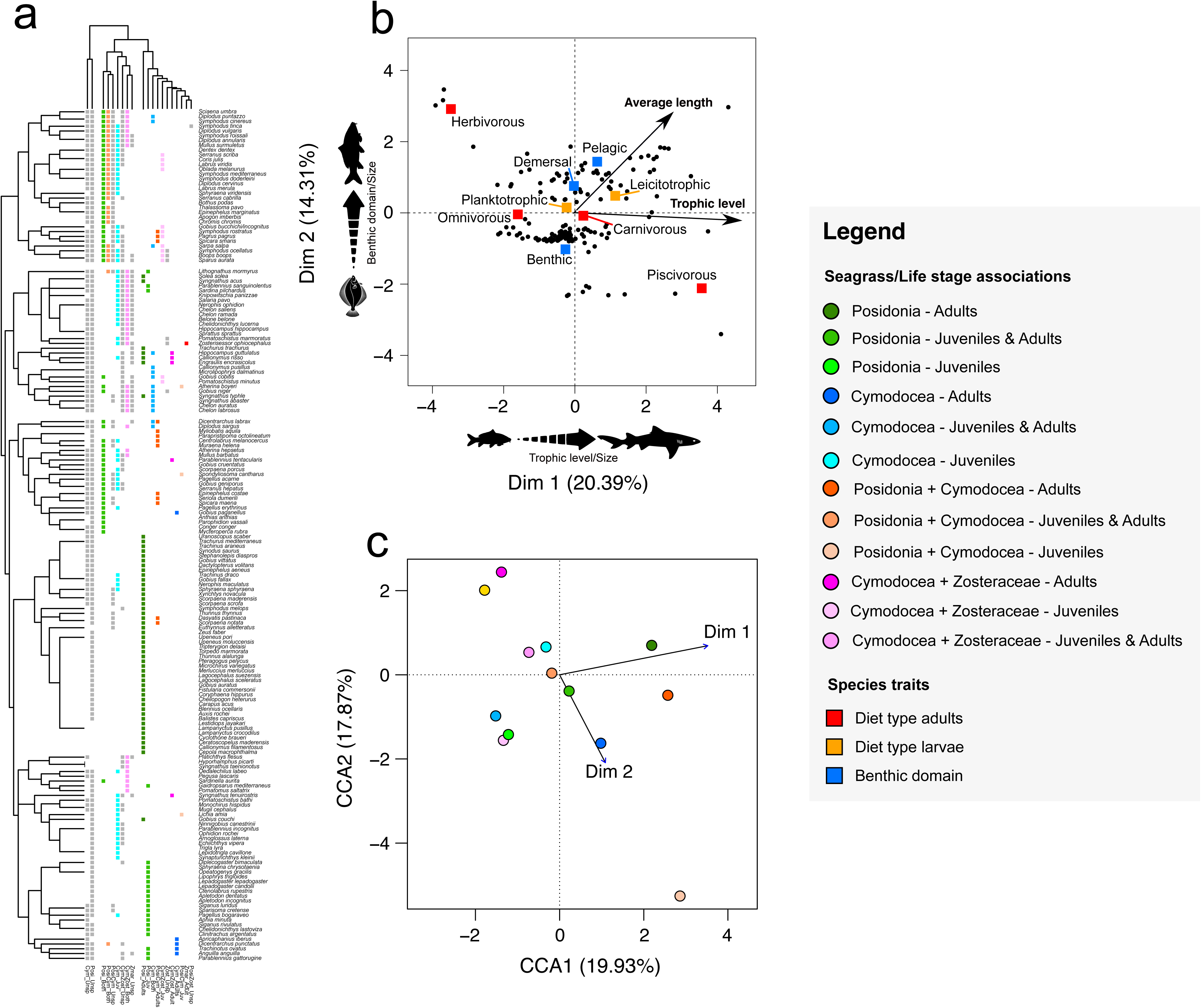
Overall picture of fish distribution, alongside their life stages and traits on different seagrass habitats. a) Heatmap showing the association between each life stage of fish reported in literature and seagrasses. Dendrograms above and alongside the heatmap represent the cluster subdivision of fish and seagrasses based on their pattern of association, using the farthest neighbour method to calculate distance between clusters in hierarchical clustering. b) Factor Analysis of Mixed Data (FAMD) showing the ordination of fish species based on functional and trophic traits. c) The first two dimensions of the FAMD (Dim 1 and Dim 2) were used in the Canonical Correspondence Analysis (CCA) to measure the association between group-level traits and seagrass habitats. Colours in the figures match the ones in the legend, with the only exception of grey-coloured squares in a), which correspond to unknown life stages of fish.

The main goal of this review was to test if species-specific traits linked to fish diet/trophic level, vertical distribution and size were associated with the presence/absence of fish species on different seagrass habitats at a given life stage. We carried out a FAMD analysis using the 175 species providing life stage data, which showed a cumulative percentage of explained variance for the first two axes of about 35% (Fig. 6b). Among continuous variables both the trophic level and, to a minor extent, the average length of the species had a strong positive association with the first dimension of the FAMD ordination (percentage of contribution 40.78% and 28.38%, respectively). Depth domain and average length mostly explained the observed pattern along the FAMD second dimension (percentage of contribution 58.54% and 13.41%, respectively). Traits related to the feeding behaviour of adults and, to a lesser extent, the vertical distribution of species, gave the highest contribution to both dimensions of the FAMD, with the first one being positively associated with large pelagic predators and the second one with low-sized species having more varying diets (Fig. 6b). When considering the pattern of ordination provided by the CCA, our results showed that adults (and to a lesser extent juveniles plus adults taken all together) inhabiting seagrass meadows of *P. oceanica* and *C. nodosa*, either pure or in combination, distributed along the first axis of the CCA (green, blue, and orange darker dots in Fig. 6c). This axis is highly correlated with the first dimension of the FAMD (*F* = 2.974, *p* = 0.003), hence suggesting that the presence of adults in these habitats is driven by both the trophic level and average length of fish species. Species in mixed seagrass habitats with Zosteraceae showed a greater dispersion along the second axis, so being loosely correlated to depth domain and size. This suggests that *P. oceanica* and *C. nodosa* habitats are significantly more populated by large carnivorous or piscivorous predators, while mixed habitats with Zosteraceae tend to host smaller species, with benthic to demersal distribution, and herbivorous to omnivorous diet.

## Discussion

In this paper we used a semi-automated procedure to recover and collate in a single database available information of fish species recorded on native seagrasses in the Mediterranean Sea, relevant for researchers working in several fields, from biodiversity conservation to fishery management. To our knowledge, our data provide the most complete information about fish associated to seagrass habitats at whole Mediterranean Sea basin scale. The development of a rigorous, structured, and iterative review process based on the PRISMA statement (Preferred Reporting Items for Systematic Reviews and Meta-Analyses) has made it possible to go beyond purely bibliometric aims, being able to exhaustively extract the scientific data needed to describe fish diversity and functional traits in key habitats (Cronin et al. 2008; Moher et al. 2009).

### Literature review and data coverage

The use of the six selected libraries for literature searching allowed recovering almost all available works, as shown by the scarce contribution provided by the *a posteriori* backward snowballing. This finding agrees with recent studies, suggesting Scopus and Web of Science as very powerful in both recalling and precision when used together (Mourão et al. 2020; Foo et al. 2021).

Although the 165 papers included in the dataset covered the entire Mediterranean Sea (Fig. 4), a geographic imbalance has been observed in the distribution of localities from which data were extracted. The western Mediterranean Sea received greater attention, with 67 sampled localities, while the Adriatic Sea had 17, and central and eastern Mediterranean had eight and nine localities, respectively. This existing gap in research has been recognized in the past and continues to persist in current literature and could be partly attributed to the less continuous distribution or even the absence of seagrasses in southern and eastern regions of the basin (Ruíz et al. 2009). However, a significant contribution to this gap comes from the disparity in study efforts between EU *vs*. non-EU countries (Ruíz et al. 2009; Chefaoui et al. 2018). Indeed, in the western Mediterranean Sea most sampling localities are located along the coasts of European countries, where *P. oceanica* beds is under protection by the European Union’s Habitats Directive (92/43/CEE) as a priority Habitat Type (*1120). Furthermore, seagrasses have all been appointed as relevant coastal bioindicators within the EU Water Framework Directive (WFD 2000/60/CE). Consequently, EU countries promoted a comprehensive knowledge of these habitats to effectively implement these Directives (Ruíz et al. 2009).

A significant portion of the studies focused on *P. oceanica*, either in pure meadows or in combination with other seagrass and seaweeds (Table 1). This is partly due to the mentioned EU legislation but the wide distribution, endemicity to Mediterranean Sea and acknowledged role in providing ecosystem services have also driven studies on various biological and ecological aspects of this seagrass (Chefaoui et al. 2018). Moreover, other studies investigated pure and mixed beds of *Cymodocea nodosa* that, although not endemic to the Mediterranean, is protected under both the Bern and OSPAR conventions being considered relevant for coastal biodiversity (Chefaoui et al. 2016). According to this sampling bias, a significant difference was observed in the number of fish species recorded, both overall and exclusive, between the species-rich *P. oceanica* and any other seagrass habitat (Table 1).

The two most frequently used sampling methods, capture and visual census, were equally applied and both effectively identified nearly all the species recorded. Investigating fish presence in seagrass meadows is challenging due to their dense multilevel canopy, so that there is not a single universally acknowledged method to efficiently survey all seagrass sub-habitats (French et al. 2021). Nevertheless, visual and capture methods complement each other, with capture being more efficient in sampling crypto-benthic species living into the canopy, while visual approaches better perform in recording planktivorous and good swimmer species (Harmelin-Vivien and Francour 1992; French et al. 2021). Accordingly, large pelagic fish as *Coryphaena hippurus*, *Euthynnys alletteratus*, *Thunnus thynnus* and *T. alalunga* were exclusively recorded through visual methods, along with other schooling and good swimmers like the blue runner *Caranx caranx* and the Mediterranean flyingfish *Cheilopogon heterurus*. On the other hand, most fish exclusively sampled through capture methods were either benthic species (e.g., *Callionymus filamentosus,* and *C. maculatus*, *Merlangius merlangus* juveniles and *U. pori*), flatfish (e.g., *Arnoglossus kessleri, Buglossidium luteum, Citharus linguatula*, *Microchirus variegatus, Platichthys flesus)* or elusive species that seek shelter under rocks and seagrass (es. *Gobius couchi*). As expected, the two methods are not totally alternative and there is no strict division among the species they efficiently sample. For instance, the marine planktivorous mesopelagic silvery lightfish *Maurolicus muelleri* was exclusively recorded by netting, while the blennid *Scartella cristata*, known to hide in empty shells or under rocks, was spotted only through visual approaches.

### Fish species in Mediterranean seagrass

The high value of Mediterranean seagrasses as marine hotspot of biodiversity has been evidenced for many macrofaunal groups, from polychaetes to amphipods (Bellisario et al. 2019; Borg et al. 2006; Buia et al. 2000; Como et al. 2008). This tenet is considered holding for fish fauna too, but there are only few papers supporting this idea by providing geographically broad data (see for example Francour 1997; Guidetti 2000; Koutsidis et al. 2020). To our knowledge, our dataset provides for the first time a comprehensive overview of fish species from native seagrass habitats across the whole Mediterranean basin.

The overall data collected confirm that seagrass habitats are biodiversity hotspots, and that the composition of fish fauna varies seasonally, although the most represented families are always the same (Labridae, Sparidae, Serranidae, Gobiidae). We found 248 fish species recorded at list twice by independent samplings (except Lessepsian species, considered even if signalled once) belonging to 75 families. This result supports the widespread postulate that seagrasses host a relevant number of fish and have therefore high importance for Mediterranean biodiversity. Indeed, considering that a total of about 650 fish species has been reported for the whole Mediterranean Sea (Coll et al. 2010), our data show that 38% of them are linked at various degrees to seagrass beds.

When considering the data at the whole basin scale, the observed pattern of records is consistent with that evidenced by single local studies. The families Labridae and Sparidae showed the highest number of records in *P. oceanica* and were the most frequent across both western and eastern Mediterranean in any sampling season, as witnessed by many studies (Deudero et al. 2008; Kalogiorou et al. 2010). Labridae was also the most frequent family recorded in *C. nodosa* and *Z. noltei*, although the composition of fish fauna in these habitats is highly variable across seasons, with necto-benthic fish prevailing in summer (Guidetti and Bussotti 2000). Highly recorded families were also Serranidae, Gobiidae, Syngnathidae and Scorpaenidae, consistently signalled as abundant in all Mediterranean seagrass communities by several studies, although their relative abundance may vary according to local conditions and human impacts (Deudero et al. 2008; Ourgaud et al. 2015). The overall data confirmed significant seasonal variations, with Serranidae as the third most recorded family after Labridae and Sparidae in summer, winter, and autumn, while Gobiidae become more frequently spotted in spring.

To investigate if the frequency of records of fish species in literature may be considered as representative of their frequency of presence in seagrass habitats, we matched their ecological and biological features with reported observations. Despite the high number of observed fish species, our data show that only a few were frequently recorded. Among the 23 most frequent species *Boops boops* and *Chromis chromis* were the most observed, both known as gregarious planktivorous species forming large aggregations on seagrass, during either reproductive season (*B. boops*) or all year long (*C. chromis*) (Deudero et al. 2008; Guidetti and Bussotti 1998). Other frequent species are known to be strictly linked to *P. oceanica* beds, as *Mullus surmuletus* and *Symphodus cinereus*, besides *Sarpa salpa* that feeds directly on *P. oceanica*, being its main herbivore consumer along with the sea urchin *Paracentrotus lividus* (Guidetti and Bussotti 2000; Máñez-Crespo et al. 2022; Prado et al. 2007). Many other species are known to be frequent and abundant in both seagrass habitats and other substrates, as *Symphodus ocellatus, S. rostratus, Serranus cabrilla, Coris julis*, and several *Diplodus* species, these latter recruiting either in seagrass or rocky substrates (Cheminée et al. 2021; Diaz-Gil et al. 2017; Garcìa-Rubies and Macpherson 1995). Some of these species are of commercial interest across the whole Mediterranean Sea (e.g., *B. boops* and *M. barbatus*) or in the Central sub-basin (e.g., *D. annularis*), further underlining the relevance of seagrass habitats in sustaining Mediterranean fisheries (FAO 2022; Jackson et al. 2015; Unsworth et al. 2018).

Our results show very low recording frequencies for most of species, whose ecology supports the idea that the low number of records really reflects their rarity in seagrass fish communities, more than representing a lack of information. Indeed, the large majority is considered rare, because of their restricted geographic range and/or low population density. For instance, the range of black-headed benny *Microlipophrys nigriceps* is limited to the western and central Mediterranean Sea and the Belotti’s goby *Gobius ater* is known from a few localities (Kovačić et al. 2022). Many uncommonly recorded species occur occasionally on seagrasses being typical of other habitats, as the killifish *Apricaphanius fasciatus*, usually living in brackish water lagoons (Mosesso et al. 2012). Habitat differences may also be associated with species’ bathymetric range, as in the case of deep-sea scaldback *Arnoglossus rueppelii* or the mesopelagic garrick *Cyclothone braueri*, one of the most abundant and frequent species of the twilight zone (Mytilineou et al. 2005; Olivar et al. 2012). Other species are pelagic and seldom occur in coastal habitats while foraging, as in the case of mackerels and tunas. Differences may also regard the substrate; this is the case of the many flatfish recorded in seagrasses although being typical of sandy bottoms as the solenette *Buglossidium luteum* or the Senegalese sole *Solea senegalensis*, that is also restricted to the western Mediterranean Sea being a Herculean migrant entered through the Gibraltar Strait (Golani et al. 2002). Some rarely recorded species are also declining throughout their whole range due to human impacts, as for endangered elasmobranchs rough ray (*Raja radula*), sandbar shark (*Carcharinus plumbeus*) and common smoothound (*Mustelus mustelus*) (Mancusi et al. 2016; Jabado et al. 2021; Rigby et al. 2021). The ecological knowledge of frequent and rare species suggests that their frequency in literature reflects their real occurrence in seagrass habitats and therefore not only confirms the high species richness of fish characterizing seagrass habitats, but also shows that only few very common species occurred, while the majority are rare and/or occasional visitors.

According to our data set, Mediterranean seagrass meadows host 15 Lessepsian species, all signalled in the ORMEF Geoportal listing the 222 exotic fish species inhabiting the Mediterranean Sea (Azzurro et al. 2022; last access August the 2^nd^, 2023). All these species are well established and quite common, although they entered the Mediterranean Sea at different times, with their first sight ranging from 1927 (*Hemiramphus far*) to 2003 (*Lagocephalus sceleratus*). All Lessepsian species have been recorded on *P. oceanica* meadows except *Sargocentron rubrum*, *Siganus luridus*, *S. rivulatus* and *Stephanolepis diaspros* that were recorded on *C. nodosa* too, and *H. far* that has been seen only a single time on *C. nodosa* (Selfati et al. 2019). The Lessepsian species spotted on seagrass in the invaded Mediterranean are mainly associated with either soft bottoms or rocky and coral reefs in their native range, with the only exception of *Sphyraena flavicauda* (Gell and Whittington 2002; Nakamura et al. 2003). This pattern is coincident with the relatively low number of records reported in seagrass according to our data, with respect to the high number of observations across the Mediterranean Sea reported in the ORMEF database (Azzurro et al. 2022).

Nevertheless, some studies have shown a high abundance of Lessepsian species in the eastern Mediterranean seagrass meadows (Kalogirou et al. 2010; Koutsidis et al. 2020). This can be attributed to the Lessepsians’ ability to expand their original niche to invade relatively empty niches, as they are often opportunistic species. Two such examples are both *Siganus* species, which are herbivorous and likely found relatively unexploited resources in the Mediterranean Sea, due to the low diversity and abundance of herbivorous fish species compared to their native ranges (Bariche et al. 2004). This allowed these species attaining high population densities, making them economically significant in terms of both landings and economic value for the eastern Mediterranean (FAO, 2022; Koutsidis et al. 2020).

### Patterns of fish diversity

Our analyses show that the high species richness of fish found on seagrass beds is far from being evenly distributed among different seagrasses. Over the 248 recorded species, Neptune seagrass habitats host the higher fish diversity, with 240 species, while 146 were observed in *C. nodosa* and 38 in *Z. marina*. These differences at the Mediterranean scale are missed when looking at the individual studies that frequently concern the comparison of seagrass habitats against bare substrates, as either rocks, gravel, or sand (e.g., Biagi et al. 1998; Franco et al. 2006; Bussotti and Guidetti 2011; La Mesa et al. 2011). Only few studies compared simultaneously and consistently fish communities associated with different seagrasses, sometimes providing contrasting results. For instance, Manent and Abella (2005) studied the fish fauna of *P. oceanica* and *C. nodosa* beds from Menorca, finding a slightly higher number of species in *C. nodosa* (16) than in *P. oceanica* (13). A similar result was found by Bonaca and Lipej (2005) within the Venice Lagoon, where the lower number of species recorded in *P. oceanica* was attributed to the very small extension of the remnant patch of this seagrass (about 1 km^2^) with respect to the area occupied by *C. nodosa*. Conversely, a comparison of juveniles’ distribution across 14 different vegetated and unvegetated habitats highlighted a higher fish species richness and abundance in *P. oceanica* with respect to *C. nodosa,* although comparable to soft and rocky substrates (Chemineé et al. 2021). A recent pioneering study investigating metazoan communities in *P. oceanica*, *C. nodosa* and the invasive *Halophila stipulacea* across the last century by using eDNA showed a higher diversity in *P. oceanica* for all examined phyla, including fish (Wesselmann et al. 2022). The Authors linked such diversity with the structural complexity of seagrass beds, known to be higher for the large *P. oceanica* than for the smaller *C. nodosa* and *H. stipulacea* (Buia et al. 2000; Hemminga and Duarte 2000).

It is worth mentioning that our dataset defined the habitats according to the main seagrass types and combinations. Detailed information on the differences among habitat patches over small scales are frequently unreported in study cases, so missing relevant information. Indeed, recent works highlighted the link between fish richness and habitat heterogeneity, especially for *P. oceanica* beds. For instance, the seascape context was found to be the main driver of Adriatic fish communities, with higher diversity associated to mosaic habitats with *P. oceanica* patches either intermingled with rocky/algal reefs and boulders or bordering sands (Zubak Čižmek et al. 2021). Accordingly, the interfaces between *P. oceanica* and other unvegetated habitats showed high species richness and abundance when looking at juvenile fish assemblages in 14 different habitats from northern Tyrrhenian Sea (Chemineé et al. 2021). Therefore, part of the high fish richness assigned to Neptune seagrass meadows could be ascribed to not reported intermix of different habitats at the local- to micro-scale, deserving specific studies so far lacking.

### Patterns of functional diversity

A main goal of this review was to investigate if traits concerning fish diet, trophic level, vertical distribution and size are related to fish species presence/absence on different seagrass habitats at a given life stage. When the presence of fish species at different life stages was associated with species-specific traits, large carnivorous or piscivorous predators resulted to be more frequently observed in *P. oceanica* and *C. nodosa* beds than in mixed habitats with Zosteraceae, which tend to host smaller species, with benthic to demersal distribution, and herbivorous to omnivorous diet.

This finding evidence a pattern of segregation between large piscivorous predators and smaller prey (including juveniles) holding at the basin scale. The dense, three-dimensional structure of seagrass may offer protection to prey with respect to bare substrates and has so generated the Seagrass Superiority Hypothesis, postulating a lower risk of predation in these habitats (Bell and Pollard 1989; Zubak et al. 2017). However, predators too may benefit from such habitats due to either the high prey density, if they are transient species, or the many hideaways if they are ambushing residents. Therefore, predation risk is not fixed for each habitat but depends largely on both the relative abundance of predators and their mode of searching and catching (Kruschel and Schultz 2020). The differential predation risk associated with different predation modes will in the end select or train prey to choose the habitat that better balances their foraging needs with the protection against the most frequent kind of predators. This is the so-called Predation Mode Hypothesis, that accounts for the role of predators in shaping fish communities and is supported by an increasing number of studies (Gilliam and Fraser 1987; Schultz et al. 2009; Fouzai et al. 2019). The results of a meta-analysis suggested that fish communities in *P. oceanic*a meadows are structured in response to the presence and abundance of piscivorous species, so indicating seagrass as a partial refuge for preys (Zubak et al. 2017). Research carried out on fish communities from vegetated and unvegetated habitats in eastern Adriatic Sea showed that prey and predators resulted negatively associated in various habitats and this negative correlation was as much stronger as predators were more aggressive. In this geographic area prey showed a positive association with *C. nodosa* and *P. oceanica*, besides unconsolidated sediments, in agreement with our analysis (Kruschel and Schultz 2020).

Our results are drawn from a high number of heterogeneous studies applying different sampling methods in different seasons, taking into consideration only seagrass habitats without bare substrates, and skipping details on predation mode. As stated in dedicated studies, local environment may play a main role in producing different patterns of prey/predator association through habitat patchiness, interspecific interactions, fishing activities and many other variables (Kruschel and Schultz 2020). Nevertheless, the signal of segregation between aggressive predators that are associated with *P. oceanica* and *C. nodosa* and their putative prey, that tend to occupy other seagrass habitats, seems to be strong enough to be recorded by our broad analyses at the whole basin scale.

## Research gaps and conclusions

This review represents a first effort to synthesize the present knowledge on fish species occurring in native seagrass habitats across the whole Mediterranean Sea, taking advantage of the high number of studies carried out so far. The database built up from the 165 papers recovered following a PRISMA modified approach provided support to several assumptions repeatedly stated in literature, but so far sustained by data from local to sub-basin scale and allowed evidencing some unexpected knowledge gaps on the highly studied fish/seagrass system.

Seagrass habitats are confirmed as key biodiversity hotspots, characterized by a high fish species richness, as already evidenced for many other taxa, with nearly 40% of the fish species recorded in the Mediterranean Sea being observed on native seagrass. However, just a minority of fish species showed high frequencies of occurrence, while the bulk was observed quite rarely on seagrass being either rare or typical of different habitats. Fish diversity is unevenly distributed among seagrasses, confirming the peculiar biodiversity hot-spot role for *P. oceanica* beds, where nearly all the fish species have been observed. However, the data so far available present a remarkable gap in terms of direct comparison among different seagrasses, since a relevant percentage of studies are cantered on the confrontation of highly different substrates as seagrasses *vs*. sediments, rocks, coralligenous, seaweeds. Recent works have been able to provide a direct confrontation of fish communities associated with different seagrasses taking advantage of the emerging eDNA techniques, so paving the way for the use of a quick and reliable approach allowing to overcome this scarcity of relevant data (He et al. 2022; Wesselmann et al. 2022).

Another key topic largely understudied is the role of habitat heterogeneity in determining fish diversity. Seagrass meadows are frequently intermingled with sandy or rocky substrates and a few recent works started to point out the relevance of habitat patchiness at the local scale in influencing fish richness. The lack of detailed studies considering habitat heterogeneity is also a limitation to understand the role of seagrasses in mediating prey/predator coexistence and in providing an effective protection as nursery. Heck and Orth (1980) attributed to complex and stratified seagrass meadows a role of refuge against predators with respect to bare substrate, hence attracting all kind of prey, including settlers and juveniles. Several studies supported this hypothesis, but some recent works started evidencing the importance of both habitat heterogeneity and most common predators’ behaviour as fundamental in determining habitat choice by prey. Aggressive piscivorous have been observed to preferentially patrol the edges among different habitats, including seagrass, so that prey species along with less aggressive predators tend to avoid heterogenous habitats. This kind of observations have generated the hypothesis that aggressive predation may be a primary constrain to fish community diversity. Although our overall data support this view at the whole basin scale, this hypothesis deserves to be tested through a higher number of diversified studies, stated its relevance for both the management of coastal resources and the implications for seagrass habitat restoration planning and implementation.

An unexpected lack of data concerned the life stage of recorded fish, that has never been reported for about one third of the species and was recorded from a minority of studies for many other species. This gap clashes with the overall acknowledged idea that seagrass beds are the habitat of choice as nursery for many fish species (Lilley and Unsworth 2014). Also, it is known that 30% to 40% of economic value of Mediterranean fisheries has been estimated to come from seagrass associated vertebrates and invertebrates. Indeed, small scale fisheries largely rely on seagrass habitats that support them both directly, being fishing grounds, and indirectly, as fish nurseries and feeding grounds (Unsworth et al. 2018). Despite this, most of recovered studies in our review were frequently oriented either at examining seasonal juveniles and adult’s occurrence in a single habitat, or at comparing highly differentiated substrates within a single geographic region, or at examining single species.

This gap is even more surprising if we consider that the degree of association between fish species and seagrasses is frequently assigned looking at the co-presence of different life stages on the same substrate (Kalogirou et al. 2012). Recently, an alternative view on the role of seagrass as nursery has been proposed, based on observations from 14 different habitats along the Mediterranean French coasts (Cheminée et al. 2021). The results obtained showed that rocky substrates were as species rich as *P. oceanica* interfaces, and that the most abundant fish species used several different habitats as nursery at various times. This agrees with global data suggesting that research is needed to fully understand the links between nursery habitats and the state of exploited fish stock (Unsworth et al. 2018). Therefore, research aimed at providing a more complete picture of the use of seagrass by fish settlers and juveniles is needed to better understand their relevance in warranting fish diversity and in sustaining Mediterranean fisheries.

## Acknowledgements

This research was carried out within a Research Training Program Scholarship of “Regione Lazio - Direzione Regionale Capitale Naturale, Parchi e Aree Protette”, which supported **AL** PhD. Research project implemented under the National Recovery and Resilience Plan (NRRP), Mission 4 Component 2 Investment 1.4 - Call for tender No. 3138 of 16 December 2021, rectified by Decree n.3175 of 18 December 2021 of Italian Ministry of University and Research funded by the European Union - Next Generation EU. Project code CN_00000033, Concession Decree No. 1034 of 17 June 2022 adopted by the Italian Ministry of University and Research, CUP J83C22000860007, Project title “National Biodiversity Future Center - NBFC”.

## Author contributions

**AL**: Methodology, investigation, formal analysis, drafted the manuscript methodology, final review; **BB**: Conceptualization, methodology, investigation, formal analysis, drafted the manuscript, final review; **RC**: Conceptualization, critical revision, drafted the manuscript, final review, investigation and supervision, resources. All authors read and approved the manuscript and agree to the submission.

## Data availability

The dataset generated and analysed during the current study is available in Figshare, https://doi.org/10.6084/m9.figshare.24100077

