## Supplementary Materials and Methods for "A review of fish diversity in Mediterranean seagrass habitats, with a focus on functional traits"

**Step 1 - Searching the libraries**

**Sub-step 1.1**

The keyword-based search was addressed to six databases: Dimension, Mendeley, ScienceDirect, Scopus, Web of Science and Wiley

The query string included the name of the Mediterranean countries, the scientific name of the Mediterranean autochthonous seagrass species belonging to the genera *Cymodocea*, *Zostera* and *Posidonia* together with the common names ‘seagrass’, ‘sea grass’, ‘eelgrass’, and the terms ‘fish’, ‘fishes’ and ‘ichthyofauna’. Through a preliminary analysis based on the method proposed by Foo and co-workers (2021) in the framework of the PRISMA approach, we added further words identifying the life history stage of fish, i.e., eggs, larvae, juveniles and adults, besides words defining the target of the observations (e.g., ‘community’ and ‘assemblage)’ and the sampling methods. This produced 69 keywords identifying the three main fields of the research objective (Mediterranean Sea, seagrass and fish fauna), linked by 68 *OR”* and three *AND”* Boolean operators.

Advanced searches were therefore carried out in each of the possible research fields (Title, Abstract, Author Keywords, Text and References) of the six analysed bibliographic databases using a different query for each library due to both their different query limits and years of coverage (Supplementary Table S1). Results have been exported as an Excel file containing the abstract, the Authors’ keywords and the title of each paper, which represents the basis of subsequent searches.

**Supplementary Table S1** - Query strings submitted to each library database.

| **Database** | **Field tags** | **Character or word limits** | **Boolean operator limits** | **Query strings** |
| --- | --- | --- | --- | --- |
| Dimensions | All Fields | None | None | (“Mediterranean Sea” OR “ Mediterranean” OR “ Alboran” OR “ Balearic” OR “ Ligurian” OR “ Tyrrhenian” OR “ Adriatic” OR “ Ionian” OR “ Aegean” OR “ Levantine” OR “ Spain” OR “ France” OR “ Monaco” OR “ Italy” OR “ Malta” OR “ Slovenia” OR “ Croatia” OR “ Bosnia-Herzegovina” OR “ Montenegro” OR “ Albania” OR “ Greece” OR “ Turkey” OR “ Cyprus” OR “ Syria” OR “ Lebanon” OR “ Israel” OR “ Palestine” OR “ Libya” OR “ Egypt” OR “ Tunisia” OR “ Algeria” OR “ Morocco”) AND (“seagrass” OR “ sea grass” OR “ eelgrass” OR “ *Posidonia oceanica*” OR “ *Cymodocea nodosa*” OR “ *Zostera marina*” OR “ *Zostera noltii*” OR “*Zostera noltei*“ OR “ phanerogam” OR “ cymodoceaceae” OR “ posidoniaceae” OR “ zosteraceae”) AND (“fish” OR “ fishes” OR “ ichthyofauna” OR “ nekton” OR “ teleostei” OR “chondrichthyes”) AND (“associated” OR “ community” OR “ assemblage” OR “ assemblages” OR “ association” OR “ communities” OR “ adult” OR “ adults” OR “ larva” OR “ larvae” OR “ egg” OR “ eggs” OR “ larval” OR “ juvenile” OR “ juveniles” OR “ nursery” OR “ visual census” OR “ sampling” OR “ capture”) |
| Mendeley | All Fields | None | None | “Mediterranean” AND “seagrass” AND “fish” AND “assemblage” |
| ScienceDirect | All Fields | 500 characters | max 8 per field | “Mediterranean” AND (“seagrass” OR “sea grass” OR “phanerogam”) AND (“fish” OR “ichthyofauna”) AND (“assemblage” OR “juvenile” OR “sampling”) |
| Scopus | All Fields | None | None | (“Mediterranean sea” OR “Mediterranean” OR “Alboran” OR “Balearic” OR “ Ligurian” OR “Tyrrhenian” OR “Adriatic” OR “Ionian” OR Aegean” OR “Levantine” OR “ Spain” OR “France” OR “Monaco” OR “Italy” OR “Malta” OR “ Slovenia” OR “Croatia” OR “Bosnia-Herzegovina” OR “ Montenegro” OR “Albania “ OR “Greece” OR “Turkey” OR “Cyprus” OR “Syria” OR “Lebanon” OR “Israel” OR “Palestine” OR “Libya” OR “Egypt” OR “ Tunisia” OR “Algeria” OR “Morocco”) AND (“seagrass” OR “sea grass” OR “eelgrass” OR “*Posidonia oceanica*” OR “*Cymodocea nodosa*” OR “*Zostera marina*” OR “*Zostera noltii*” OR “*Zostera noltei*” OR “phanerogam” OR “cymodoceaceae” OR “posidoniaceae” OR “zosteraceae”) AND (“fish” OR “fishes” OR “ichthyofauna” OR “nekton” OR “teleostei” OR “chondrichthyes”) AND (“associated” OR “community” OR “assemblage” OR “assemblages” OR “association” OR “communities” OR “adult” OR “adults” OR “larva” OR “larvae” OR “egg” OR “eggs” OR “larval” OR “juvenile” OR “juveniles” OR “nursery” OR “visual census” OR “sampling” OR “capture”) |
| WebOfScience | All Fields | 50 terms for “All Fields” | None | (“Mediterranean” OR “Alboran” OR “Balearic” OR “Ligurian” OR “Tyrrhenian” OR “Adriatic” OR “Ionian” OR “Aegean” OR “Levantine”) AND (“seagrass” OR “sea grass” OR “eelgrass” OR “*Posidonia oceanica*” OR “*Cymodocea nodosa*” OR “*Zostera marina*” OR “*Zostera noltii*” OR “*Zostera noltei*” OR “phanerogam” OR “cymodoceaceae” OR “posidoniaceae” OR “zosteraceae”) AND (“fish” OR “fishes” OR “ichthyofauna” OR “nekton” OR “teleostei” OR “chondrichthyes”) AND (“associated” OR “community” OR “assemblage” OR “assemblages” OR “association” OR “communities” OR “visual census” OR “nursery” OR “sampling” OR “capture”) |
| Wiley Online Library | All fields | None | None | (“Mediterranean sea” OR “Mediterranean” OR “Alboran” OR “Balearic” OR “ Ligurian” OR “Tyrrhenian” OR “Adriatic” OR “Ionian” OR Aegean” OR “Levantine” OR “ Spain” OR “France” OR “Monaco” OR “Italy” OR “Malta” OR “ Slovenia” OR “Croatia” OR “Bosnia-Herzegovina” OR “ Montenegro” OR “Albania “ OR “Greece” OR “Turkey” OR “Cyprus” OR “Syria” OR “Lebanon” OR “Israel” OR “Palestine” OR “Libya” OR “Egypt” OR “ Tunisia” OR “Algeria” OR “Morocco”) AND (“seagrass” OR “sea grass” OR “eelgrass” OR “*Posidonia oceanica*” OR “*Cymodocea nodosa*” OR “*Zostera marina*” OR “*Zostera noltii*” OR “*Zostera noltei*” OR “phanerogam” OR “cymodoceaceae” OR “posidoniaceae” OR “zosteraceae”) AND (“fish” OR “fishes” OR “ichthyofauna” OR “nekton” OR “teleostei” OR “chondrichthyes”) AND (“associated” OR “community” OR “assemblage” OR “assemblages” OR “association” OR “communities” OR “adult” OR “adults” OR “larva” OR “larvae” OR “egg” OR “eggs” OR “larval” OR “juvenile” OR “juveniles” OR “nursery” OR “visual census” OR “sampling” OR “capture”) |

**Sub-step 1.2 - Removing duplicates and selecting key-terms**

After collecting the search results, we removed from the paper list records having partial searchable information (i.e., literature without abstract) and duplicate papers, by using both Microsoft Excel and the function ‘removes.duplicates’ in the package *litsearchr* of R (Grames et al. 2019; R Core Team 2022).

We subsequently performed a new search for a higher, more specific and exhaustive number of key‑terms (193) related to the four key-subjects (hereafter groups) of the research: Group S - Seagrass, 47 key-terms related to the seagrasses and habitat targets; “Group F - Fish, 52 key-terms related to fish biology; “Group A - Assemblage, 42 general key-terms related to either the sampling methods or fish assemblages; “Group G – Geographic locality, 53 key-terms related to the Mediterranean basin and geographic localities (Supplementary Table S2).

**Supplementary Table S2** - Key-terms subdivided in the four subject groups.

| **Key-subjects** | **Key-terms** |
| --- | --- |
| S - SEAGRASS | “Angiosperm”; “ Bed”; “ Bedform”; “ Bedforms”; “ Beds”; “ Canopies”; “ Canopy”; “ Cover”; “ Cymodocea”; “ Cymodoceaceae”; “ Eelgrass”; “ Grass”; “ Grasses”; “ Intertidal”; “ Leaves”; “ Macrophyte”; “ Marina”; “ Matte”; “ Meadow”; “ Meadows”; “ Monocots”; “ Monocotyledons”; “ MPAS”; “ Nanozostera”; “ Neptune”; “ Nodosa”; “ Noltei”; “ Noltii”; “ Oceanica”; “ Phanerogam”; “ Phanerogams”; “ Plant”; “ Plants”; “ Posidonia”; “ Posidoniaceae”; “ Protected”; “ Rhizome”; “ Seagrass”; “ Seagrasses”; “ Substratum”; “ Subtidal”; “ Tapeweed”; “ Tidal”; “ Vegetated”; “ Vegetation”; “ Zostera”; “ Zosteraceae” |
| F - FISH | “Adult”; “ Adults”; “ Assemblage”; “ Assemblages”; “ Benthic”; “ Benthos”; “ Breeding”; “Chondrichthyes”; “ Communities”; “ Community”; “ Demersal”; “ Egg”; “ Eggs”; “ Fauna”; “ Feeding”; “ Fish”; “ Fishes”; “ Grazer”; “ Grazers”; “ Ichthyofauna”; “ Ichthyofaunal”; “ Juvenile”; “ Juveniles”; “ Larva”; “ Larvae”; “ Larval”; “ Lifespan”; “ Necton”; “ Nectonic”; “ Nekton”; “ Nektonic”; “ Nurseries”; “ Nursery”; “ Pisces”; “ Piscivore”; “ Piscivorous”; “ Piscivory”; “ Recruit”; “ Recruitment”; “ Recruits”; “ Settlement”; “ Settler”; “ Settlers”; “ Spawn”; “ Spawner” ; “ Spawners”; “ Spawning”; “ Stock”; “ Teleost”; “ Teleostei”; “ Teleosts” |
| A - ASSEMBLAGE | “Capture”; “ Captures”; “ Catch”; “ Catches”; “ Census”; “ Censused”; “ Censuses”; “ Coast”; “ Coastal”; “ Count”; “ Counts”; “ Discarded”; “ Discards”; “ Fished”; “ Fisheries”; “ Fishery”; “ Fishing”; “ Littoral”; “ Net”; “ Nets”; “ Occurrence”; “ Occurrences”; “ Recapture”; “ Recaptured”; “ Residence”; “ Residency”; “ Sample”; “ Sampled”; “ Samples”; “ Sampling”; “ Samplings”; “ Seine”; “ Seiners”; “ Telemetry”; “ Tracking”; “ Transect”; “ Transects”; “ Trawl”; “ Trawling”; “ Trawls”; “ Video”; “ Visual” |
| G - GEOGRAPHY | “Adriatic”; “ Aegean”; “ Albania”; “ Albanian”; “ Alboran”; “ Algeria”; “ Algerian”; “ Balearic”; “ Bosnia”; “ Bosnian”; “ Croatia”; “ Croatian”; “ Cypriot”; “ Cyprus”; “ Egypt”; “ Egyptian”; “ France”; “ French”; “ Greece”; “ Greek”; “ Ionian”; “ Israel”; “ Israeli”; “ Italian”; “ Italy”; “ Lebanese”; “ Lebanon”; “ Levantine”; “ Libya”; “ Libyan”; “ Ligurian”; “ Malta”; “ Maltese”; “ Mediterranean”; “ Monacan”; “ Monaco”; “ Montenegrin”; “ Montenegro”; “ Moroccan”; “ Morocco”; “ Palestine”; “ Palestinian”; “ Slovenia”; “ Slovenian”; “ Spain”; “ Spanish”; “ Syria”; “ Syrian”; “ Tunisia”; “ Tunisian”; “ Turkey”; “ Turkish”; “ Tyrrhenian” |

**Step 2 – Scoring and filtering papers**

**Sub-step 2.1**

In the second step, we detected the presence/absence of the key-terms above mentioned in Title, Abstract and Authors’ keywords of each search result using the 'str_detect' function in the package stringr of R (Wickham 2019).

To quickly identify papers containing relevant information, we defined a “Relevance Score (RS) built by assigning a relative weight to each of the four key-subject groups. We assigned a maximum value of one (100%) to those papers dealing with (hence containing key-terms related to) all the four key-subject groups, and modulated the score attributed to each group according to the level of relevance for our research objective, as follows:

1) 1/3 of the score was assigned if the paper contained at least one key-term belonging to Group S- Seagrass, i.e., Group S = 0.333;
2) 1/3 of the score was assigned if the paper contained at least one key-term belonging to Group F- Fish, i.e., Group F = 0.333;
3) 2/9 of the score were assigned if the paper contained at least one key-term belonging to Group A - Assemblage, i.e., Group A = 0.222;
4) 1/9 of the score was assigned if the paper contained at least one key-term belonging to Group G - Geographic locality, i.e., Group G = 0.111.

The Relevance Score (RS) was therefore based on presence/absence of the key-terms and summed up the values attributed to all key-terms found in each paper, ranging from 0 to 1. Papers with a RS ≥ 0.667 contain either both key-terms from high relevant key-subject groups (i. e., groups S or F) or one of them in combination with both key-terms from the other groups (A, G). Based on this, papers were subdivided in “High Relevance Papers” (HRP, RS ≥ 0.667) and “Low Relevance Papers“ (LRP, RS <0.667) (Supplementary Table S3).

**Supplementary Table S3** - Relevance Score values obtained from the various possible combinations of key-terms *per* key-subject group found in the abstract and title of the papers. High Relevance Score combinations are evidenced in bold.

| **Group S** | **Group F** | **Group A** | **Group G** | **Relevance Score Values** |
| --- | --- | --- | --- | --- |
| **0.333** | **0.333** | **0.222** | **0.111** | **1** |
| **0.333** | **0.333** | **0.222** | **0** | **0.889** |
| **0.333** | **0.333** | **0** | **0.111** | **0.778** |
| **0.333** | **0.333** | **0** | **0** | **0.667** |
| **0** | **0.333** | **0.222** | **0.111** | **0.667** |
| **0.333** | **0** | **0.222** | **0.111** | **0.667** |
| 0.333 | 0 | 0.222 | 0 | 0.556 |
| 0 | 0.333 | 0.222 | 0 | 0.556 |
| 0.333 | 0 | 0 | 0.111 | 0.444 |
| 0 | 0.333 | 0 | 0.111 | 0.444 |
| 0 | 0 | 0.222 | 0.111 | 0.333 |
| 0.333 | 0 | 0 | 0 | 0.333 |
| 0 | 0.333 | 0 | 0 | 0.333 |
| 0 | 0 | 0.222 | 0 | 0.222 |
| 0 | 0 | 0 | 0.111 | 0.111 |

**Sub-step 2.2-– Filter #1**

We performed a sequential filtering to keep only those papers containing available information to build up the final dataset. In this first filtering step, we analysed the overall number of key‑terms belonging to each of the four key‑subject groups (i.e., not only their presence/absence as in the previous step) to set a threshold value based on the representativeness of each of the four key‑subject groups in each paper.

To this end, we counted for each paper the overall number of key‑terms belonging to each key‑subject group and summed them up in four combinations: i) SFAG = all four groups (i.e., all the key‑terms retrieved); ii) SFG = the sum of the key‑terms belonging to the subject groups related to fish, seagrass and the Mediterranean Sea; iii) SFA = the sum of the key‑terms belonging to subjects groups related to fish, seagrass and assemblages; iv) SF = the sum of the key‑terms belonging to subject groups related to fish and seagrass. This allowed associating four different values to each paper, one for every combination (*n*). For each *n* we then counted the corresponding number of HRPs and LRPs (as reported in Supplementary Table S4). This allowed us to: 1) calculate the difference among HRPs and LRPs for each *n* (ΔRP); 2) find the highest ΔRP for each *n*, corresponding to the higher key‑terms presence in HRPs with respect to LRP; 3) set the number of key‑terms corresponding to the highest ΔRP for each *n* as a threshold (Supplementary Table S5); 4) filter out the papers having a number of key‑terms lower than the threshold values for even a single combination.

**Supplementary Table S4** - Filter 1. Summary of ΔRP values obtained for the combination of the four key-‑subject groups SFAG, SFG, SFA and SF. n: the total number of Key-terms retrieved; LRP and HRP: total number of papers containing n key-terms; ΔRP: difference between LRP and HRP papers for each n recorded. The maximum difference observed is evidenced in red bold characters.

| **SFAG** | | | | **SFG** | | | | **SFA** | | | | **SF** | | | |
| --- | --- | --- | --- | --- | --- | --- | --- | --- | --- | --- | --- | --- | --- | --- | --- |
| **n** | **LRP** | **HRP** | **ΔRP** | **n** | **LRP** | **HRP** | **ΔRP** | **n** | **LRP** | **HRP** | **ΔRP** | **n** | **LRP** | **HRP** | **ΔRP** |
| 0 | 1737 | 0 | -1737 | 0 | 4143 | 0 | -4143 | 0 | 1935 | 0 | -1935 | 0 | 5016 | 0 | -5016 |
| 1 | 1845 | 0 | -1845 | 1 | 3392 | 0 | -3392 | 1 | 2090 | 0 | -2090 | 1 | 3323 | 581 | -2742 |
| 2 | 2132 | 174 | -1958 | 2 | 2476 | 795 | -1681 | 2 | 2248 | 325 | -1923 | 2 | 2284 | 1250 | -1034 |
| 3 | 1863 | 448 | -1415 | 3 | 1720 | 1239 | -481 | 3 | 1843 | 676 | -1167 | 3 | 1545 | 1521 | -24 |
| 4 | 1798 | 632 | -1166 | 4 | 1441 | 1349 | -92 | 4 | 1725 | 868 | -857 | 4 | 1308 | 1442 | 134 |
| 5 | 1444 | 845 | -599 | 5 | 999 | 1392 | 393 | 5 | 1379 | 1042 | -337 | 5 | 918 | 1438 | 520 |
| 6 | 1290 | 981 | -309 | 6 | 800 | 1285 | 485 | 6 | 1198 | 1137 | -61 | 6 | 740 | 1289 | 549 |
| 7 | 1029 | 1007 | -22 | **7** | **583** | **1271** | **688** | 7 | 971 | 1141 | 170 | **7** | **543** | **1107** | **564** |
| 8 | 915 | 1015 | 100 | 8 | 514 | 1161 | 647 | 8 | 856 | 1097 | 241 | 8 | 480 | 1041 | 561 |
| 9 | 715 | 1090 | 375 | 9 | 375 | 1024 | 649 | 9 | 684 | 1028 | 344 | 9 | 378 | 873 | 495 |
| 10 | 617 | 1036 | 419 | 10 | 332 | 927 | 595 | 10 | 583 | 1002 | 419 | 10 | 312 | 799 | 487 |
| 11 | 502 | 971 | 469 | 11 | 242 | 802 | 560 | **11** | **472** | **967** | **495** | 11 | 224 | 692 | 468 |
| 12 | 417 | 902 | 485 | 12 | 218 | 680 | 462 | 12 | 407 | 832 | 425 | 12 | 211 | 633 | 422 |
| **13** | **306** | **862** | **556** | 13 | 179 | 625 | 446 | 13 | 282 | 776 | 494 | 13 | 167 | 528 | 361 |
| 14 | 280 | 757 | 477 | 14 | 145 | 538 | 393 | 14 | 261 | 677 | 416 | 14 | 140 | 457 | 317 |
| 15 | 208 | 714 | 506 | 15 | 104 | 500 | 396 | 15 | 201 | 608 | 407 | 15 | 103 | 426 | 323 |
| 16 | 191 | 616 | 425 | 16 | 105 | 402 | 297 | 16 | 182 | 572 | 390 | 16 | 100 | 340 | 240 |
| 17 | 189 | 532 | 343 | 17 | 71 | 360 | 289 | 17 | 183 | 473 | 290 | 17 | 67 | 333 | 266 |
| 18 | 121 | 507 | 386 | 18 | 70 | 314 | 244 | 18 | 117 | 440 | 323 | 18 | 66 | 274 | 208 |
| 19 | 111 | 459 | 348 | 19 | 47 | 268 | 221 | 19 | 105 | 398 | 293 | 19 | 41 | 244 | 203 |
| 20 | 83 | 386 | 303 | 20 | 35 | 256 | 221 | 20 | 83 | 324 | 241 | 20 | 35 | 208 | 173 |
| 21 | 66 | 343 | 277 | 21 | 30 | 214 | 184 | 21 | 67 | 337 | 270 | 21 | 32 | 173 | 141 |
| 22 | 52 | 315 | 263 | 22 | 23 | 180 | 157 | 22 | 50 | 271 | 221 | 22 | 22 | 149 | 127 |
| 23 | 61 | 273 | 212 | 23 | 18 | 155 | 137 | 23 | 56 | 249 | 193 | 23 | 13 | 129 | 116 |
| 24 | 34 | 249 | 215 | 24 | 14 | 133 | 119 | 24 | 31 | 194 | 163 | 24 | 11 | 106 | 95 |
| 25 | 28 | 214 | 186 | 25 | 8 | 106 | 98 | 25 | 30 | 181 | 151 | 25 | 10 | 73 | 63 |
| 26 | 20 | 196 | 176 | 26 | 9 | 77 | 68 | 26 | 19 | 161 | 142 | 26 | 8 | 67 | 59 |
| 27 | 6 | 160 | 154 | 27 | 1 | 94 | 93 | 27 | 6 | 122 | 116 | 27 | 1 | 63 | 62 |
| 28 | 7 | 147 | 140 | 28 | 3 | 71 | 68 | 28 | 7 | 107 | 100 | 28 | 3 | 48 | 45 |
| 29 | 8 | 119 | 111 | 29 | 6 | 49 | 43 | 29 | 6 | 92 | 86 | 29 | 4 | 47 | 43 |
| 30 | 10 | 94 | 84 | 30 | 2 | 32 | 30 | 30 | 9 | 65 | 56 | 30 | 1 | 19 | 18 |
| 31 | 8 | 75 | 67 | 31 | 2 | 36 | 34 | 31 | 8 | 56 | 48 | 31 | 2 | 23 | 21 |
| 32 | 7 | 58 | 51 | 32 | 2 | 19 | 17 | 32 | 6 | 47 | 41 | 32 | 1 | 17 | 16 |
| 33 | 4 | 55 | 51 | 33 | 0 | 21 | 21 | 33 | 4 | 26 | 22 | 33 | 0 | 16 | 16 |
| 34 | 2 | 36 | 34 | 34 | 1 | 12 | 11 | 34 | 2 | 35 | 33 | 34 | 1 | 6 | 5 |
| 35 | 0 | 23 | 23 | 35 | 0 | 10 | 10 | 35 | 0 | 25 | 25 | 35 | 0 | 5 | 5 |
| 36 | 3 | 31 | 28 | 36 | 0 | 10 | 10 | 36 | 3 | 14 | 11 | 36 | 0 | 7 | 7 |
| 37 | 0 | 20 | 20 | 37 | 0 | 7 | 7 | 37 | 0 | 10 | 10 | 37 | 0 | 2 | 2 |
| 38 | 0 | 15 | 15 | 38 | 0 | 8 | 8 | 38 | 0 | 15 | 15 | 38 | 0 | 2 | 2 |
| 39 | 0 | 20 | 20 | 39 | 0 | 6 | 6 | 39 | 0 | 14 | 14 | 39 | 0 | 3 | 3 |
| 40 | 0 | 15 | 15 | 40 | 0 | 2 | 2 | 40 | 0 | 11 | 11 | 40 | 0 | 3 | 3 |
| 41 | 0 | 12 | 12 | 41 | 0 | 3 | 3 | 41 | 0 | 5 | 5 | 41 | 0 | 2 | 2 |
| 42 | 0 | 7 | 7 | 42 | 0 | 2 | 2 | 42 | 0 | 2 | 2 | 43 | 0 | 1 | 1 |
| 43 | 0 | 5 | 5 | 43 | 0 | 1 | 1 | 43 | 0 | 5 | 5 | 45 | 0 | 2 | 2 |
| 44 | 0 | 7 | 7 | 45 | 0 | 2 | 2 | 44 | 0 | 2 | 2 | 46 | 0 | 2 | 2 |
| 45 | 0 | 3 | 3 | 46 | 0 | 2 | 2 | 45 | 0 | 1 | 1 |  |  |  |  |
| 46 | 0 | 3 | 3 | 51 | 0 | 1 | 1 | 46 | 0 | 2 | 2 |  |  |  |  |
| 47 | 1 | 1 | 0 |  |  |  |  | 47 | 1 | 1 | 0 |  |  |  |  |
| 49 | 0 | 4 | 4 |  |  |  |  | 48 | 0 | 2 | 2 |  |  |  |  |
| 50 | 0 | 1 | 1 |  |  |  |  | 49 | 0 | 2 | 2 |  |  |  |  |
| 51 | 0 | 2 | 2 |  |  |  |  | 50 | 0 | 1 | 1 |  |  |  |  |
| 52 | 0 | 1 | 1 |  |  |  |  | 51 | 0 | 2 | 2 |  |  |  |  |
| 55 | 0 | 2 | 2 |  |  |  |  | 54 | 0 | 1 | 1 |  |  |  |  |
| 60 | 0 | 1 | 1 |  |  |  |  |  |  |  |  |  |  |  |  |

According to the results rereported in Table S4, we retained the papers containing at least 13, 7, 11 and 7 key-terms (Supplementary Table S5).

**Supplementary Table S5** - Key-terms combinations listed with the threshold values (n), the total number of LRPs and HRPs characterized by that n, and the corresponding ΔRP.

|  | **n** | **LRP** | **HRP** | **ΔRP** |
| --- | --- | --- | --- | --- |
| ALL | 13 | 306 | 862 | 556 |
| FSG | 7 | 583 | 1271 | 688 |
| FSA | 11 | 472 | 967 | 495 |
| FS | 7 | 543 | 1107 | 564 |

**Sub-step 2.2 -– Filter 2**

After their use in filter 1, we discarded LRPs papers as they contain scanty information for our research objective (i.e., RS < 0.667).

**Sub-step 2.3 – Filter 3**

A further filtering was implemented to exclude the papers that, although containing most of the 193 key-terms, may be empty of relevant information. To this end, we selected 35 “Negative Keywords” (NK) from LRPs, by means of a Document-Term Matrix (DTM). Formally, a DTM is a word frequency table where columns are words and rows are documents, with entries representing the frequency of retrieval of any specific word in each document. We used the function ‘TermDocumentMatrix’ from the *tm* package of R (Feinerer and Hornik 2023), and the total number of NK was obtained from the average number of Authors’ keywords. We vectorized the Authors’ keywords of each paper removing any punctuation and special character and converted the text to lower case using the package *tm* of R before DTM. The selected NK therefore identified keywords frequently present in research providing irrelevant information, because carried out outside the Mediterranean basin or not focusing on seagrass and/or fish communities although mentioning them, for instance for comparison purposes. (Supplementary List 1)

**Supplementary List S1** - List of the 35 Negative Keywords NK. We used these keywords to filter out the papers that contain even just one of these words in their Most Frequent Keywords (see M&M): “America”; “Anthozoa”; “Aquaculture”; “Australia”; “Bioaccumulation”; “Botany”; “Brazil”; “Carbon”; “Chemical”; “Chemistry”; “China”; “Coral”; “Crustacea”; “Estuarine”; “Estuary”; “Eutrophication”; “Gene”; “Genetic”; “Isotope”; “Isotopes”; “Mercury”; “Metal”; “Metals”; “Mexico”; “Nitrogen”; “Photosynthesis”; “Physiological”; “Physiology”; “Pollutant”; “Pollutants”; “Pollution”; “Portugal”; “River”; “Sediment”; “Sediments”

We obtained the “Most Frequent Keywords” (MFK) of HRPs identifying the recurring and relevant keywords of each paper. To this end, in papers featuring the Authors’ keywords we retained them as MFK. In papers where keywords were absent, we extracted the 40 terms most frequently reported in the title and abstract of each paper using a value of frequency calculated again through a Document-Term Matrix (DTM) and obtaining the number of most frequent words equal to the average number of the MFK present in the papers featuring the Authors’ keywords.

We filtered the list of papers by removing: i) all papers containing one or more NK within their MFK; ii) all papers with a Key-Terms/MFK ratio lower than 0.2, i.e. all papers that contain a Key-Terms/MFK ratio lower than MFK/(n key-subject groups + 1) ratio ; iii) all papers that do not have any keyword related to Group S (seagrass) or F (fish) in their MFK.

**Results**

Fish records

In our dataset, after manual data extraction from the papers and data validation, a total of 248 fish species were detected within the 10 habitat combinations, belonging to 75 families. For each species, scientific name was checked in World Register of Marine Species (WoRMS) using the “match taxa” tool provided in the WoRMS website, last access on August 10, 2023

**Supplementary Table S6.** - List of the 248 species recovered from literature subdivided by family, with their frequency of observation. Colour codes correspond to Fig. S1 (see below): white (rare), grey (intermediate), black (frequent). Lessepsian species are highlighted in red.

| **Family** | **Scientific Name** | **Frequency** |
| --- | --- | --- |
| Alosidae | *Alosa fallax* |  |
| Alosidae | *Sardina pilchardus* |  |
| Ammodytidae | *Gymnammodytes cicerelus* |  |
| Anguillidae | *Anguilla anguilla* |  |
| Apogonidae | *Apogon imberbis* |  |
| Atherinidae | *Atherina boyeri* |  |
| Atherinidae | *Atherina hepsetus* |  |
| Atherinidae | *Atherina presbyter* |  |
| Balistidae | *Balistes capriscus* |  |
| Belonidae | *Belone belone* |  |
| Blenniidae | *Aidablennius sphynx* |  |
| Blenniidae | *Blennius ocellaris* |  |
| Blenniidae | *Coryphoblennius galerita* |  |
| Blenniidae | *Lipophrys trigloides* |  |
| Blenniidae | *Microlipophrys canevae* |  |
| Blenniidae | *Microlipophrys dalmatinus* |  |
| Blenniidae | *Microlipophrys nigriceps* |  |
| Blenniidae | *Parablennius gattorugine* |  |
| Blenniidae | *Parablennius incognitus* |  |
| Blenniidae | *Parablennius pilicornis* |  |
| Blenniidae | *Parablennius rouxi* |  |
| Blenniidae | *Parablennius sanguinolentus* |  |
| Blenniidae | *Parablennius tentacularis* |  |
| Blenniidae | *Parablennius zvonimiri* |  |
| Blenniidae | *Salaria pavo* |  |
| Blenniidae | *Scartella cristata* |  |
| Bothidae | *Arnoglossus kessleri* |  |
| Bothidae | *Arnoglossus laterna* |  |
| Bothidae | *Arnoglossus rueppelii* |  |
| Bothidae | *Arnoglossus thori* |  |
| Bothidae | *Bothus podas* |  |
| Callionymidae | *Callionymus filamentosus* |  |
| Callionymidae | *Callionymus maculatus* |  |
| Callionymidae | *Callionymus pusillus* |  |
| Callionymidae | *Callionymus risso* |  |
| Carangidae | *Caranx crysos* |  |
| Carangidae | *Lichia amia* |  |
| Carangidae | *Pseudocaranx dentex* |  |
| Carangidae | *Seriola dumerili* |  |
| Carangidae | *Trachinotus ovatus* |  |
| Carangidae | *Trachurus mediterraneus* |  |
| Carangidae | *Trachurus trachurus* |  |
| Carapidae | *Carapus acus* |  |
| Carcharhinidae | *Carcharhinus plumbeus* |  |
| Cepolidae | *Cepola macrophthalma* |  |
| Citharidae | *Citharus linguatula* |  |
| Clinidae | *Clinitrachus argentatus* |  |
| Clupeidae | *Sprattus sprattus* |  |
| Congridae | *Ariosoma balearicum* |  |
| Congridae | *Conger conger* |  |
| Coryphaenidae | *Coryphaena hippurus* |  |
| Cyprinodontidae | *Aphanius fasciatus* |  |
| Cyprinodontidae | *Apricaphanius iberus* |  |
| Dactylopteridae | *Dactylopterus volitans* |  |
| Dasyatidae | *Dasyatis pastinaca* |  |
| Dorosomatidae | *Sardinella aurita* |  |
| Engraulidae | *Engraulis encrasicolus* |  |
| Exocoetidae | *Cheilopogon heterurus* |  |
| Fistulariidae | *Fistularia commersonii* |  |
| Gadidae | *Merlangius merlangus* |  |
| Gobiesocidae | *Apletodon dentatus* |  |
| Gobiesocidae | *Apletodon incognitus* |  |
| Gobiesocidae | *Diplecogaster bimaculata* |  |
| Gobiesocidae | *Lepadogaster candolii* |  |
| Gobiesocidae | *Lepadogaster lepadogaster* |  |
| Gobiesocidae | *Opeatogenys gracilis* |  |
| Gobiidae | *Aphia minuta* |  |
| Gobiidae | *Buenia affinis* |  |
| Gobiidae | *Deltentosteus collonianus* |  |
| Gobiidae | *Deltentosteus quadrimaculatus* |  |
| Gobiidae | *Gobius ater* |  |
| Gobiidae | *Gobius auratus* |  |
| Gobiidae | *Gobius bucchichi/incognitus* |  |
| Gobiidae | *Gobius cobitis* |  |
| Gobiidae | *Gobius couchi* |  |
| Gobiidae | *Gobius cruentatus* |  |
| Gobiidae | *Gobius fallax* |  |
| Gobiidae | *Gobius geniporus* |  |
| Gobiidae | *Gobius niger* |  |
| Gobiidae | *Gobius paganellus* |  |
| Gobiidae | *Gobius vittatus* |  |
| Gobiidae | *Gobius xanthocephalus* |  |
| Gobiidae | *Knipowitschia panizzae* |  |
| Gobiidae | *Ninnigobius canestrinii* |  |
| Gobiidae | *Odondebuenia balearica* |  |
| Gobiidae | *Pomatoschistus bathi* |  |
| Gobiidae | *Pomatoschistus marmoratus* |  |
| Gobiidae | *Pomatoschistus minutus* |  |
| Gobiidae | *Pomatoschistus quagga* |  |
| Gobiidae | *Pseudaphya ferreri* |  |
| Gobiidae | *Thorogobius ephippiatus* |  |
| Gobiidae | *Thorogobius macrolepis* |  |
| Gobiidae | *Zebrus zebrus* |  |
| Gobiidae | *Zosterisessor ophiocephalus* |  |
| Gonostomatidae | *Cyclothone braueri* |  |
| Haemulidae | *Parapristipoma octolineatum* |  |
| Haemulidae | *Pomadasys incisus* |  |
| Hemiramphidae | *Hemiramphus far* |  |
| Hemiramphidae | *Hyporhamphus picarti* |  |
| Holocentridae | *Sargocentron rubrum* |  |
| Labridae | *Centrolabrus melanocercus* |  |
| Labridae | *Coris julis* |  |
| Labridae | *Ctenolabrus rupestris* |  |
| Labridae | *Labrus merula* |  |
| Labridae | *Labrus mixtus* |  |
| Labridae | *Labrus viridis* |  |
| Labridae | *Pteragogus pelycus* |  |
| Labridae | *Symphodus cinereus* |  |
| Labridae | *Symphodus doderleini* |  |
| Labridae | *Symphodus mediterraneus* |  |
| Labridae | *Symphodus melops* |  |
| Labridae | *Symphodus ocellatus* |  |
| Labridae | *Symphodus roissali* |  |
| Labridae | *Symphodus rostratus* |  |
| Labridae | *Symphodus tinca* |  |
| Labridae | *Thalassoma pavo* |  |
| Labridae | *Xyrichtys novacula* |  |
| Lophiidae | *Lophius piscatorius* |  |
| Lotidae | *Gaidropsarus mediterraneus* |  |
| Lotidae | *Gaidropsarus vulgaris* |  |
| Merlucciidae | *Merluccius merluccius* |  |
| Monacanthidae | *Stephanolepis diaspros* |  |
| Moronidae | *Dicentrarchus labrax* |  |
| Moronidae | *Dicentrarchus punctatus* |  |
| Mugilidae | *Chelon auratus* |  |
| Mugilidae | *Chelon labrosus* |  |
| Mugilidae | *Chelon ramada* |  |
| Mugilidae | *Chelon saliens* |  |
| Mugilidae | *Mugil cephalus* |  |
| Mugilidae | *Oedalechilus labeo* |  |
| Mullidae | *Mullus barbatus* |  |
| Mullidae | *Mullus surmuletus* |  |
| Mullidae | *Upeneus moluccensis* |  |
| Mullidae | *Upeneus pori* |  |
| Muraenidae | *Muraena helena* |  |
| Myctophidae | *Ceratoscopelus maderensis* |  |
| Myctophidae | *Lampanyctus crocodilus* |  |
| Myctophidae | *Lampanyctus pusillus* |  |
| Myctophidae | *Myctophum punctatum* |  |
| Myliobatidae | *Myliobatis aquila* |  |
| Ophichthidae | *Echelus myrus* |  |
| Ophichthidae | *Ophisurus serpens* |  |
| Ophidiidae | *Ophidion barbatum* |  |
| Ophidiidae | *Ophidion rochei* |  |
| Ophidiidae | *Parophidion vassali* |  |
| Paralepididae | *Lestidiops jayakari* |  |
| Phycidae | *Phycis phycis* |  |
| Pleuronectidae | *Platichthys flesus* |  |
| Pomacentridae | *Chromis chromis* |  |
| Pomatomidae | *Pomatomus saltatrix* |  |
| Rajidae | *Raja asterias* |  |
| Rajidae | *Raja clavata* |  |
| Rajidae | *Raja radula* |  |
| Scaridae | *Sparisoma cretense* |  |
| Sciaenidae | *Sciaena umbra* |  |
| Sciaenidae | *Umbrina cirrosa* |  |
| Scomberesocidae | *Scomberesox saurus* |  |
| Scombridae | *Auxis rochei* |  |
| Scombridae | *Euthynnus alletteratus* |  |
| Scombridae | *Sarda sarda* |  |
| Scombridae | *Scomber colias* |  |
| Scombridae | *Thunnus alalunga* |  |
| Scombridae | *Thunnus thynnus* |  |
| Scophthalmidae | *Scophthalmus maximus* |  |
| Scophthalmidae | *Scophthalmus rhombus* |  |
| Scorpaenidae | *Pterois miles* |  |
| Scorpaenidae | *Scorpaena maderensis* |  |
| Scorpaenidae | *Scorpaena notata* |  |
| Scorpaenidae | *Scorpaena porcus* |  |
| Scorpaenidae | *Scorpaena scrofa* |  |
| Scyliorhinidae | *Scyliorhinus canicula* |  |
| Scyliorhinidae | *Scyliorhinus stellaris* |  |
| Serranidae | *Anthias anthias* |  |
| Serranidae | *Epinephelus aeneus* |  |
| Serranidae | *Epinephelus caninus* |  |
| Serranidae | *Epinephelus costae* |  |
| Serranidae | *Epinephelus marginatus* |  |
| Serranidae | *Mycteroperca rubra* |  |
| Serranidae | *Serranus atricauda* |  |
| Serranidae | *Serranus cabrilla* |  |
| Serranidae | *Serranus hepatus* |  |
| Serranidae | *Serranus scriba* |  |
| Siganidae | *Siganus luridus* |  |
| Siganidae | *Siganus rivulatus* |  |
| Soleidae | *Buglossidium luteum* |  |
| Soleidae | *Microchirus ocellatus* |  |
| Soleidae | *Microchirus variegatus* |  |
| Soleidae | *Monochirus hispidus* |  |
| Soleidae | *Pegusa impar* |  |
| Soleidae | *Pegusa lascaris* |  |
| Soleidae | *Solea senegalensis* |  |
| Soleidae | *Solea solea* |  |
| Soleidae | *Synapturichthys kleinii* |  |
| Sparidae | *Boops boops* |  |
| Sparidae | *Dentex dentex* |  |
| Sparidae | *Dentex gibbosus* |  |
| Sparidae | *Diplodus annularis* |  |
| Sparidae | *Diplodus cervinus* |  |
| Sparidae | *Diplodus puntazzo* |  |
| Sparidae | *Diplodus sargus* |  |
| Sparidae | *Diplodus vulgaris* |  |
| Sparidae | *Lithognathus mormyrus* |  |
| Sparidae | *Oblada melanurus* |  |
| Sparidae | *Pagellus acarne* |  |
| Sparidae | *Pagellus bogaraveo* |  |
| Sparidae | *Pagellus erythrinus* |  |
| Sparidae | *Pagrus pagrus* |  |
| Sparidae | *Sarpa salpa* |  |
| Sparidae | *Sparus aurata* |  |
| Sparidae | *Spicara maena* |  |
| Sparidae | *Spicara smaris* |  |
| Sparidae | *Spondyliosoma cantharus* |  |
| Sphyraenidae | *Sphyraena chrysotaenia* |  |
| Sphyraenidae | *Sphyraena flavicauda* |  |
| Sphyraenidae | *Sphyraena sphyraena* |  |
| Sphyraenidae | *Sphyraena viridensis* |  |
| Sternoptychidae | *Maurolicus muelleri* |  |
| Syngnathidae | *Hippocampus guttulatus* |  |
| Syngnathidae | *Hippocampus hippocampus* |  |
| Syngnathidae | *Nerophis maculatus* |  |
| Syngnathidae | *Nerophis ophidion* |  |
| Syngnathidae | *Syngnathus abaster* |  |
| Syngnathidae | *Syngnathus acus* |  |
| Syngnathidae | *Syngnathus taenionotus* |  |
| Syngnathidae | *Syngnathus tenuirostris* |  |
| Syngnathidae | *Syngnathus typhle* |  |
| Synodontidae | *Synodus saurus* |  |
| Tetraodontidae | *Lagocephalus sceleratus* |  |
| Tetraodontidae | *Lagocephalus suezensis* |  |
| Torpedinidae | *Torpedo marmorata* |  |
| Torpedinidae | *Torpedo torpedo* |  |
| Trachinidae | *Echiichthys vipera* |  |
| Trachinidae | *Trachinus araneus* |  |
| Trachinidae | *Trachinus draco* |  |
| Trachinidae | *Trachinus radiatus* |  |
| Triakidae | *Mustelus mustelus* |  |
| Triglidae | *Chelidonichthys cuculus* |  |
| Triglidae | *Chelidonichthys lastoviza* |  |
| Triglidae | *Chelidonichthys lucerna* |  |
| Triglidae | *Chelidonichthys obscurus* |  |
| Triglidae | *Eutrigla gurnardus* |  |
| Triglidae | *Lepidotrigla cavillone* |  |
| Triglidae | *Trigla lyra* |  |
| Tripterygiidae | *Tripterygion delaisi* |  |
| Tripterygiidae | *Tripterygion melanurus* |  |
| Tripterygiidae | *Tripterygion tripteronotum* |  |
| Uranoscopidae | *Uranoscopus scaber* |  |
| Zeidae | *Zeus faber* |  |

**Slope Analysis**

The Akaike Information Criterion (AIC, Burnham and Anderson 1998) identified the piecewise regression as the best model in explaining the relationship between the number of records for each of the 248 fish species reported in literature and the number of localities where they were observed, AIC_piecewise_ = 1743.752, AIC_linear_ = 1933.885 and ΔAIC = 190.133. The piecewise regression improved the adjusted R-squared (from 0.951 to 0.978), identifying two main breakpoints at ca. 17 and 55 on the x-axis, corresponding to a significant change in slope (Fig. S1). Computations were performed by using the *segmented* package of R (Muggeo 2008).

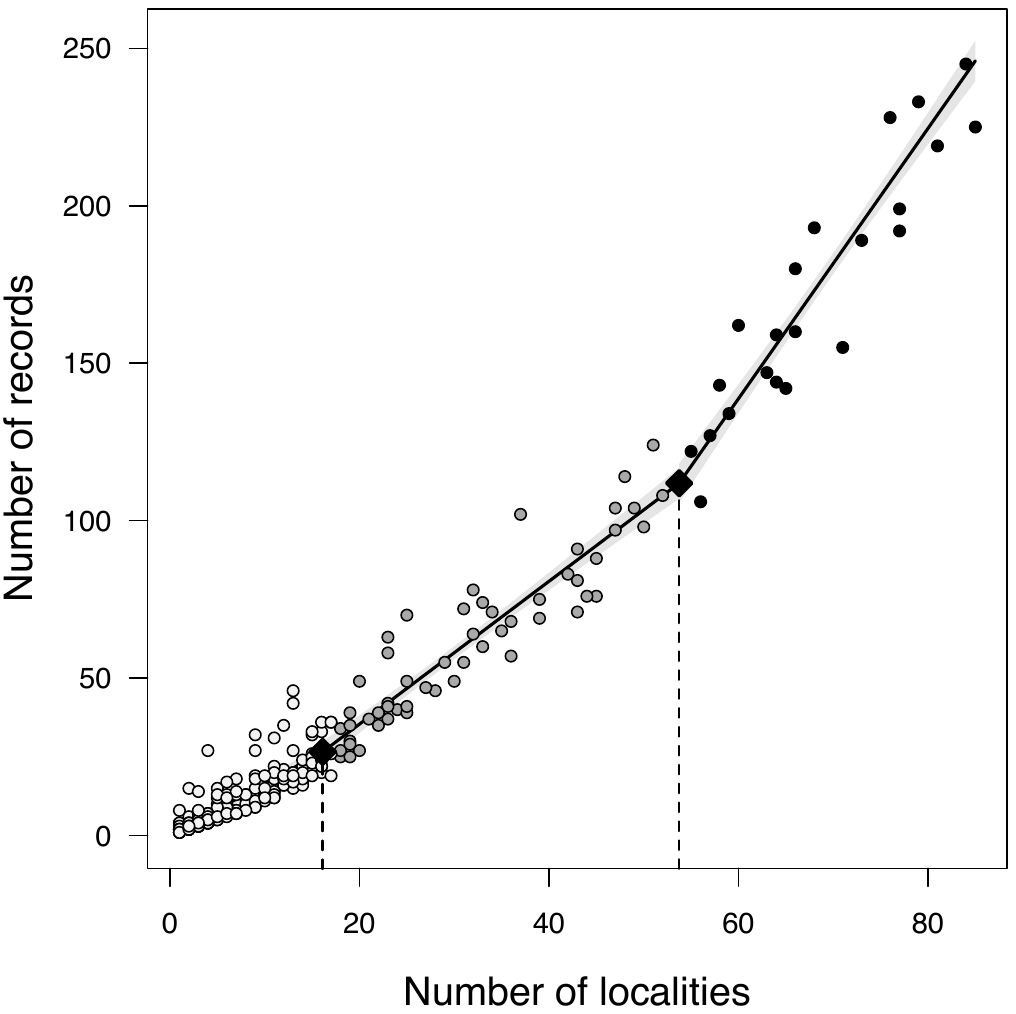

**Supplementary Figure S1** - The curve slope analysis identifies two main breakpoints (black diamonds) corresponding to a significant change in the slope of regression and the consequent subdivision of species (coloured circles) in three main groups: white for rare species, grey for intermediate frequency, black for common species (see Supplementary Table S6).
